# A persistent invasive phenotype in post-hypoxic tumor cells is revealed by novel fate-mapping and computational modeling

**DOI:** 10.1101/2020.12.30.424757

**Authors:** Heber L. Rocha, Inês Godet, Furkan Kurtoglu, John Metzcar, Kali Konstantinopoulos, Soumitra Bhoyar, Daniele M. Gilkes, Paul Macklin

**Affiliations:** Department of Intelligent Systems Engineering, Indiana University, Bloomington, IN 47408, USA; Department of Oncology, The Sidney Kimmel Comprehensive Cancer Center, The Johns Hopkins University School of Medicine, Baltimore, MD 21231, USA; Department of Chemical and Biomolecular Engineering, The Johns Hopkins University, Baltimore, MD 21218, USA; Department of Informatics, Indiana University, Bloomington, IN 47408, USA; Cellular and Molecular Medicine Program, The Johns Hopkins University School of Medicine, Baltimore, MD 21231, USA

## Abstract

Hypoxia is a critical factor in solid tumors that has been associated with cancer progression and aggressiveness. We recently developed a hypoxia-fate mapping system that allowed the tracing of post-hypoxic cells within a tumor for the first time. This novel approach, based on an oxygen-dependent fluorescent switch, made the investigation of the post-hypoxic phenotype possible. The system allowed us to measure key biological features such as oxygen distribution, cell proliferation and migration. Using this data, we developed a computational model to investigate the motility and phenotypic persistence of hypoxic and post-hypoxic cells during tumor progression. The behavior of hypoxic and post-hypoxic cells was defined by phenotypic persistence time, cell movement bias and the fraction of cells that respond to an enhanced migratory stimulus. Our studies revealed that post-hypoxic cells have an enhanced persistent migratory phenotype that promotes the formation of invasive structures (“plumes”) expanding towards the oxygenated tumor regions. This work combined advanced cell tracking and imaging techniques with mathematical modeling, and revealed for the first time that a persistent invasive migratory phenotype that develops under hypoxic conditions enhances their escape into non-hypoxic tumor regions to invade the surrounding tissue.

## INTRODUCTION

Intratumoral hypoxia, or oxygen (O_2_) deprivation, is associated with an increased risk of metastasis, treatment failure and worse patient outcome (Gilkes and Semenza, 2013;Muz et al., 2015). Hypoxia occurs in 90% of solid tumors as a result of rapid cancer cell proliferation and inefficient vasculature (Michiels, 2008). Studies using an Eppendorf electrode demonstrated that the median partial pressure of oxygen (pO_2_) in solid tumors in patients with breast cancer was 10 mmHg (1.3% O_2_) compared to normal breast tissue whose median pO_2_ was 65 mmHg (8.6% O_2_) (Vaupel et al., 2007).

The cellular response to hypoxia is driven by the hypoxia-inducible factors 1 and 2 (HIF-1 and HIF-2). The alpha subunits of HIF-1 and HIF-2 are subject to proteosomal degradation under well-oxygenated conditions. Under hypoxia, HIF-1*α* and HIF-2*α* are stabilized, translocate to the nucleus, and heterodimerize with HIF-1*β* (Berra et al., 2001). HIF-1 and HIF-2 heterodimers transcriptionally regulate gene expression when O_2_ is limited. Hypoxia has been reported to regulate the expression of more than a thousand genes (Ye et al., 2018), the majority of which are induced by HIFs.

Current methods to detect hypoxia within a tumor are limited to distinguishing cells that are hypoxic at the time of measurement. Consequently, upon reoxygenation, the hypoxic stimulus is lost, and the ultimate fate of the post-hypoxic cells is unknown. To overcome this obstacle, we recently developed a novel hypoxia fate-mapping system that uses a Loxp-Cre approach to drive a permanent fluorescent switch from DsRed to GFP expression once a cell experiences hypoxia. Using this strategy, our preliminary data demonstrated that cells exposed to hypoxia in the primary tumor are 5 times more efficient at forming lung metastasis in mouse models of breast cancer (Godet et al., 2019). Moreover, as we monitored tumor progression in an orthotopic mouse model during a time-course experiment, we visualized the appearance of GFP+ tumor regions lining the necrotic core of the tumor that eventually progress to larger GFP+ areas.

Although these observations were insightful, we could not fully characterize the dynamic cellular changes that occur in response to the O_2_ gradients within the tumor. Intravital imaging would allow us to acquire real-time dynamics in select areas of the tumor; however, it is limited to short time intervals where the animal is immobilized on the microscope stage and only small tumor volumes can be analyzed. Therefore, the goal of our current work is to establish a computational model based on *in vivo* and *ex vivo* data that captures the spatio-temporal dynamics of hypoxic cells at large length scales, and use it to explore potential hypotheses driving post-hypoxic cell behavior.

There is a large body of work on mathematical modeling of cell migration in the context of tumor progression (Anderson et al., 2000; Prahl and Odde, 2018; Rangarajan and Zaman, 2008; Stonko et al.,2015). To date, many mathematical models have included hypoxia when modeling the dynamics of tumor growth (Alarcón et al., 2004; Lima et al., 2014; Macklin et al., 2012; Macnamara et al., 2020; Meaney et al., 2019; Rocha et al., 2018).Macklin et al. (2012) used a model for ductal carcinoma *in situ* where cells exposed to hypoxia could not recover to their non-hypoxic state, and immediately became necrotic. In Lima et al. (2014), hypoxic cells released pro-angiogenic factors such as VEGF into the tumor microenvironment, which stimulated endothelial cells to promote angiogenesis and partially relieved hypoxia. A model combining hypoxia-activated prodrugs (HAP) with radiation therapy was developed by Meaney et al. (2019). The migratory advantage of hypoxic cells was modeled by Macnamara et al.(2020). However, modeling both hypoxic and post-hypoxic cells has not been attempted due to the lack of experimental data required to build such model.

Here, we have the unique opportunity to gather biological data from cells within different tumor regions: necrotic and viable. The tumor cells have distinct phenotypic states according to O_2_ distribution: normoxic cells localize in oxygenated tumor regions (DsRed+), hypoxic cells localize adjacent to the necrotic core (GFP+), post-hypoxic cells localize in oxygenated regions (GFP+) after migration, and non-viable cells localize in the necrotic core (Figure 1A). Experimental data gathered from this set-up were used to establish a multiscale continuum-discrete model within the PhysiCell simulation framework (Ghaffarizadeh et al., 2018). *In vivo* O_2_ measurements performed in tumors using an optical probe defined the O_2_ gradient. To obtain motility measurements, we used tumor-isolated cancer cells cultured in an *ex vivo* 3D spheroid model. Using the approximate Bayesian computation (ABC) technique (Beaumont et al., 2002), we calibrated key parameters to model cell motility derived from biological experiments, particularly speed and movement bias (Figure 1B). Proliferation was assessed using Ki67 staining, which marks cells in active and inactive phases of the cell cycle (Figure 1C).

**Figure 1.**
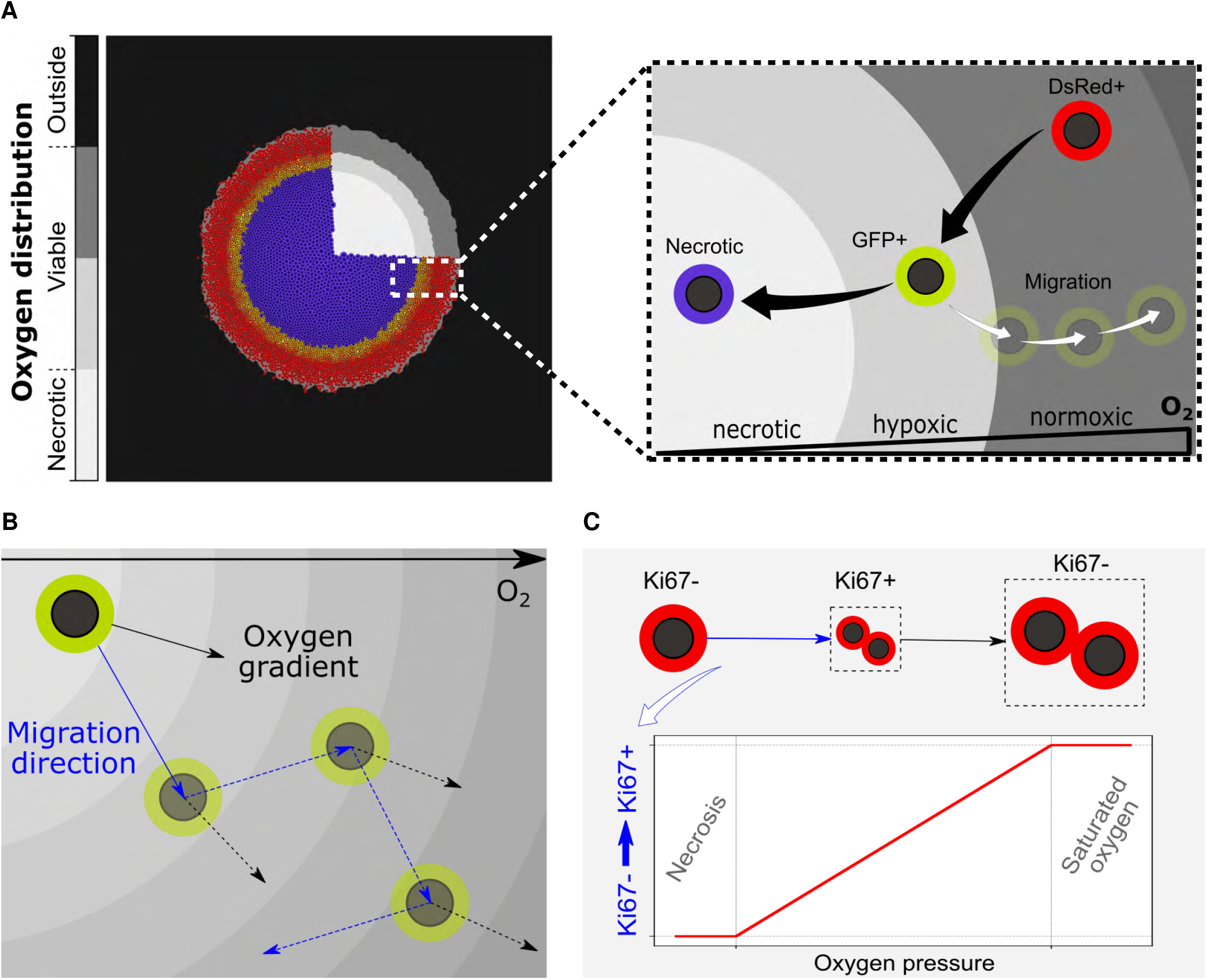
Overview of the computational model. (A) The O_2_ distribution in the microenvironment defines the viable and necrotic regions of the tumor (left). Phenotypic changes defined by the concentration of O_2_ and permanent switch to GFP fluorescence (right). (B) Random migration with a directional bias along the O_2_ gradient. (C) Scheme of the cell cycle, where the transition rate from Ki67-(non-cycling) to Ki67+ (cycling) scales with O_2_ availability.

To design a representative model of tumor progression, we defined the following parameters to characterize the GFP+ cells: phenotypic persistence time (*T_p_*), bias of the cell migration (*b*\*), and fraction of cells that can respond to a migration stimulus (*F_r_*). Phenotypic persistence time is defined as the persistence of the migratory response once the cell leaves the hypoxic region. The motility bias illustrates the tendency of cell migration to align with the O_2_ gradient. Lastly, the fraction of responders represents the number of GFP+ cells that experience an enhanced migration when compared to the remaining GFP+ population, as suggested by our biological experiments. By testing a range of these parameters, we explored different scenarios predicted by our model and compared them to the *in vivo* observations from tissue sections of the primary tumor both in 2D and 3D. A qualitative comparison with the in vivo output demonstrates that our model accurately recapitulates the biological observations of GFP+ invasive structures.

This work combines the novel hypoxia fate-mapping system with computational modeling to gain new biological insights on the dynamical behavior of post-hypoxic cancer cells. The hypoxia fate-mapping allows us to visualize novel invasive tumor structures, while the computational model allows us to explore the cell behavioral hypotheses and dynamics that can and cannot recapitulate the novel invasive structures. We then “close the loop”using targeted experimental imaging to test the simulation hypotheses. In particular, this integrative approach will show that a persistent post-hypoxic phenotype is needed to explain “plumes” of GFP+ cells in well-oxygenated tumor regions. The work demonstrates how the marriage of novel imaging and computational modeling can drive advances in understanding hypoxia-driven cancer invasion.

## RESULTS

### Fate-mapping intratumoral hypoxia enables characterization of a hypoxic/post-hypoxic phenotype

We recently developed a novel approach to fate-map hypoxic cells *in vivo* during tumor progression in murine models of breast cancer (Godet et al., 2019). The model was developed by delivering two lentiviral vectors to the human metastatic breast cancer cell line, MDA-MB-231. The first vector contains a constitutively active promoter and encodes an expression cassette for a red fluorescent protein (DsRed) flanked by tandem loxP sites and followed by a gene encoding a green fluorescent protein (GFP). The second vector contains four hypoxia responsive elements (HREs) that transcriptionally regulate the expression of a Cre gene which was altered by adding an oxygen-dependent degradation domain (ODD). Consequently, under hypoxia, Cre promotes the cleavage of the DsRed gene leading to the permanent expression of GFP (Figure 2A, 2B).

**Figure 2.**
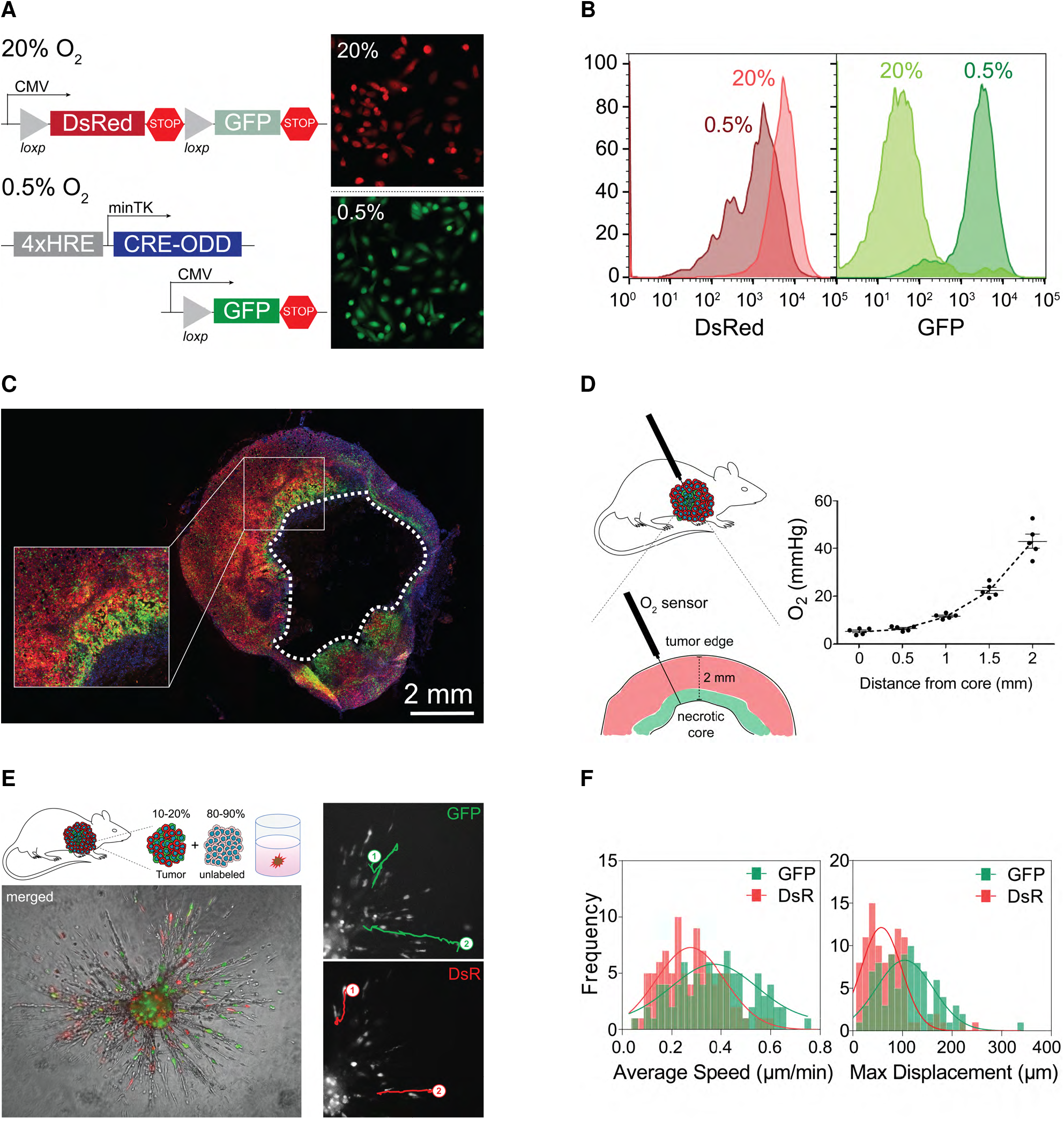
Fate-mapping intratumoral hypoxia. (A,B) Lentiviral vectors used to generate the hypoxia fate-mapping system. Fluorescent images (A) and flow cytometry analysis (B) of the MDA-MB-231 hypoxia fate-mapping cells following exposure to 20% or 0.5% O_2_ for 5 days. (C) Fluorescent images of the full cross-section of an orthotopic tumor derived from MDA-MB-231 hypoxia fate-mapping cells. (D) O_2_ measurements performed using a fixed-needle micro-probe are displayed as a function of distance from the tumor core. (E,F) Representative spheroid used to track migrating tumor-derived GFP+ or DsRed+ cells (E) and corresponding average speed and maximum displacement of individual cells (F) fit by Gaussian non-linear regression.

To investigate the emergence and progression of hypoxia in primary breast tumors, hypoxia fatemapping MDA-MB-231 cells were implanted in the mammary fat pad of NSG mice. Tumors were harvested, cryo-sectioned, mounted and imaged following 15, 20, 35 and 40 days of tumor growth (Figure 2C, Supplementary figure 1a). GFP+ cells were localized to peri-necrotic regions of the tumor, and GFP+ areas within the tumor increased over time (Godet et al., 2019). We also observed the leading edge of GFP+ tumor areas invading into DsRed+ tumor regions (Figure 2C).

In order to build a mathematical model of our *in vivo* system, we conducted experiments to define key parameters, such as the spatial distribution of O_2_ in MDA-MB-231-derived orthotopic tumors and the proliferative capacity of GFP+ and DsRed+ cells. Using an optical detection probe, O_2_ measurements were recorded at 0.5 mm depth increments within the 2 mm viable rim. The maximum recorded pO_2_ was 43-46 mmHg at the periphery of the tumor, while in the perinecrotic region, the pO_2_ dropped to 6 mmHg (Figure 2D). To assess the proliferative capacity of DsRed+ and GFP+ cells in tumors, we immunofluorescently labeled tumor tissue sections with Ki67, which marks cells in active phases of the cell cycle (Bruno and Darzynkiewicz, 1992). Image analysis revealed that overall, 50 to 55% of cancer cells were Ki67+, which is in line with previously reported Ki67 indexes in MDA-MB-231 derived tumors (Warin et al., 2010). Moreover, there was no significant difference between Ki67 expression in DsRed+ and GFP+ cells (Supplementary figure 1b,c), suggesting that proliferation is not distinct between the two cell populations.

Given previous findings (Ju et al., 2020, 2017; Lewis et al., 2016) that hypoxia enhances cell migration and invasion, we investigated the migratory-invasive behavior of tumor-derived cells in a spheroid model. In our studies, we isolated GFP+ and DsRed+ cells from primary orthotopic mouse tumors and used them to generate spheroids containing 10-20% tumor-derived cells and 80-90% unlabeled type cells. Using live cell imaging, we tracked individual cells migrating within the spheroid as well as invading into the adjacent collagen matrix (Figure 2E). GFP+ cells displayed a higher mean speed and maximum displacement (*s* = 0.38 *μm*/*min* and *d_max_* = 111.11 *μm*) compared to DsRed+ cells (*s* = 0.28 *μm*/*min* and *d_max_* = 68.34 *μm*) (Figure 2F). The observations suggest that even after reoxygenation, GFP+ cells are more motile than DsRed+ cells.

### Biological observations calibrate cellular motility in hypoxia computational model

We calibrated our model based on the previously described cell motility experiments using tumor-derived cells. To extract the displacement profile of these cells, we simulated the random migration tracks of 100 cells seeded at the same initial position. We modeled each cell’s migration as independent to focus on the distribution of individual cell tracks. We set the motility persistence time (*τ*) to 15 minutes to match the experimental sampling frequency, and we calibrated the speed (*s*) and the bias (*b*) for DsRed+ and GFP+ cells. By applying Bayesian inference with ABC methodology, in particular the LF-MCMC method (Marjoram et al., 2003), we obtained the posterior distribution of (*b,s*) conditioned to the experimental data *y_o_* for the simulated displacement of the DsRed+ and GFP+ cells (Figure 3A).

**Figure 3.**
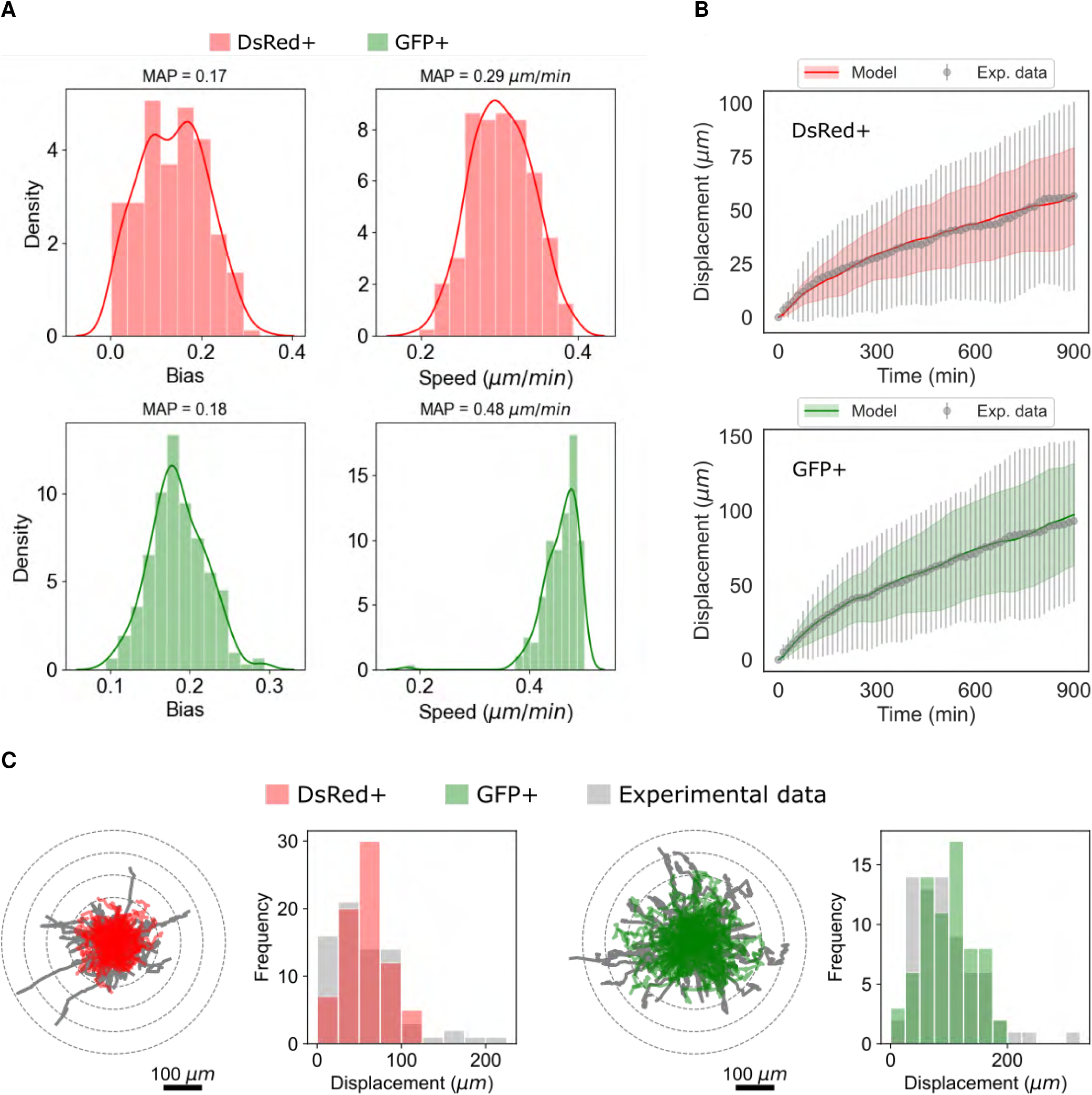
Calibration of cell migration in the model. (A) Frequency histograms of the posterior distributions for *b* (migration bias) and *s* (migration speed) for DsRed+ cells (labeled red) and GFP+ cells (labeled green) using the LF-MCMC method. (B) Longitudinal changes in displacement (mean and standard deviation) obtained by simulating the model with the MAP (*ℓ*_2_ norm of the residuals between the data average and the simulated values are approximately 12.9 *μm*). (C) Trajectories of all cells (74 cells DsRed+ and 69 cells GFP+) over 15 hours and frequency histogram of the displacement, in which the gray graphs are the biological data, and the red and green graphs are associated with the DsRed + and GFP + cells, respectively.

According to the distributions obtained, we extracted the maximum *a posteriori* probability (MAP) of speed and bias, and we recomputed the simulated cell tracks using these estimated parameters. Finally, we compared the simulated displacements and trajectories with the experimental data and verified that our model was effective at recapitulating the biological observations (Figure 3B, 3C). Moreover, the average displacement obtained by the model closely approximated the observational data for both cells types, specifically, the residual for DsRed+ and GFP+ cells was 12.92 *μm* and 12.86 *μm*, respectively. The cell trajectories have a similar profile when compared to the experimental data; however, trajectories with greater displacement were observed in the experimental data, which might suggest that a sub-population of GFP+ cells have a stronger response to migration stimuli.

### Computational model captures diverse spatial-temporal dynamic patterns

To study the effects of the enhanced migratory phenotype of the GFP+ cells on the spatiotemporal progression of the tumor without the confounding complexities of *in vivo* models, we designed a computational model for virtual thought experiments (Macklin, 2017, 2019). In our virtual system, we simulated a 3 mm square (2D) or cube (3D) with an initial tumor comprised entirely of DsRed+ cells. We arranged the cells at the center of the domain in a packed disc with a radius 0.25 mm. We then applied the parameters calibrated in the previous section, along with other parameters defined according to literature, or estimated as specified in Supplementary Table 1. We used this virtual system to assess the impact of varying the simulation rules that correspond to hypotheses on cell phenotypic responses to hypoxic conditions. Because there is uncertainty in how long a GFP+ cell can maintain its migratory phenotype after leaving a region of hypoxia, we defined a parameter for phenotypic persistence time (*T_p_*) and analyzed the impact of its variation on the dynamics of tumor evolution. We first assessed the impact of phenotypic persistence when hypoxic (GFP+) cells had the calibrated migrational bias (*b* = 0.1718) under three sets of hypotheses:

1. **No phenotypic persistence** (*T_p_* =0 *h*): This represents the hypothesis that GFP+ cells do not show phenotypic persistence; they immediately resume a normoxic, non-migratory phenotype upon leaving hypoxic tissue regions.
2. **Transient phenotypic persistence** (0 <*T_p_* < ∞): This represents the case where a hypoxic cell maintains its hypoxic migratory phenotype for a certain duration (*T_p_*) after leaving hypoxic regions. We fixed *T_p_* = 50 *h*.
3. **Permanent phenotypic persistence** (*T_p_* = ∞): This represents the scenario where hypoxic cells maintain their migratory phenotype permanently.

In this first virtual experiment, we found that increasing *T_p_* promoted the mixing of DsRed+ and GFP+ cells along the hypoxic boundary region and in the viable rim. Furthermore, no GFP+ cells were able to escape through the viable rim to invade the surrounding tissue (Supplementary Figure 2).

Because this does not match our *in vivo* observations where GFP+ cells frequently invade the surrounding tissues and seed distant micrometastases, we surmise that additional behavioral rules are needed for the GFP+ cells. Phenotypic persistence in the absence of strong directional cues for migration cannot drive invasion in the computational model system.

We next varied the motility bias (*b*) of the GFP+ cells and set *T_p_* to 0 to assess whether changes in the randomness of migration alone could drive invasion of GFP+ cells. We evaluated the changes associated with either partially Brownian (0 < *b* < 1), or completely polarized (*b* = 1) movement along the oxygen gradient ∇σ. We found that increasing *b* caused the overall tumor to spread more quickly, allowing hypoxic cells to escape prior to necrotic death. However, none of these cells were able to completely escape the viable tumor region to begin invasion. Thus, increased directionality of cell migration alone cannot explain tissue invasion (Supplementary Figure 3).

We then investigated whether the combination of increased migrational polarization (increased migration bias *b*) and phenotypic persistence (*T_p_* > 0) could lead to tissue invasion by GFP+ cells. As noted, increasing migrational bias (*b* = 0.5) with no phenotypic persistence (*T_p_* = 0) did not lead to GFP+ invasion beyond the tumor edge. Instead, the tumor grew radially, and it was composed of three homogeneous layers of necrotic, GFP+ and DsRed+ cells (Figure 4A). However, when we introduced transient (*T_p_* = 50 *h*) or permanent (*T_p_* = ∞) phenotypic persistence, we found that GFP+ cells formed “plume” structures that invaded into the well-oxygenated DsRed+ regions of the tumor (Figure 4B, 4C). For sufficiently long phenotypic persistence times, we found that GFP+ cells eventually migrated through the entire DsRed+ tumor region to invade the surrounding tissue (see *t* =130 *h* in Figure 4B (*T_p_* = 50 *h*) and Figure 4C (*T_p_* = ∞)). Thus, a persistence hypoxic migratory phenotype – when coupled with sufficiently directed migration – is sufficient to drive GFP+ cell invasion through the non-hypoxic DsRed+ tumor regions and into the surrounding tissue.

**Figure 4.**
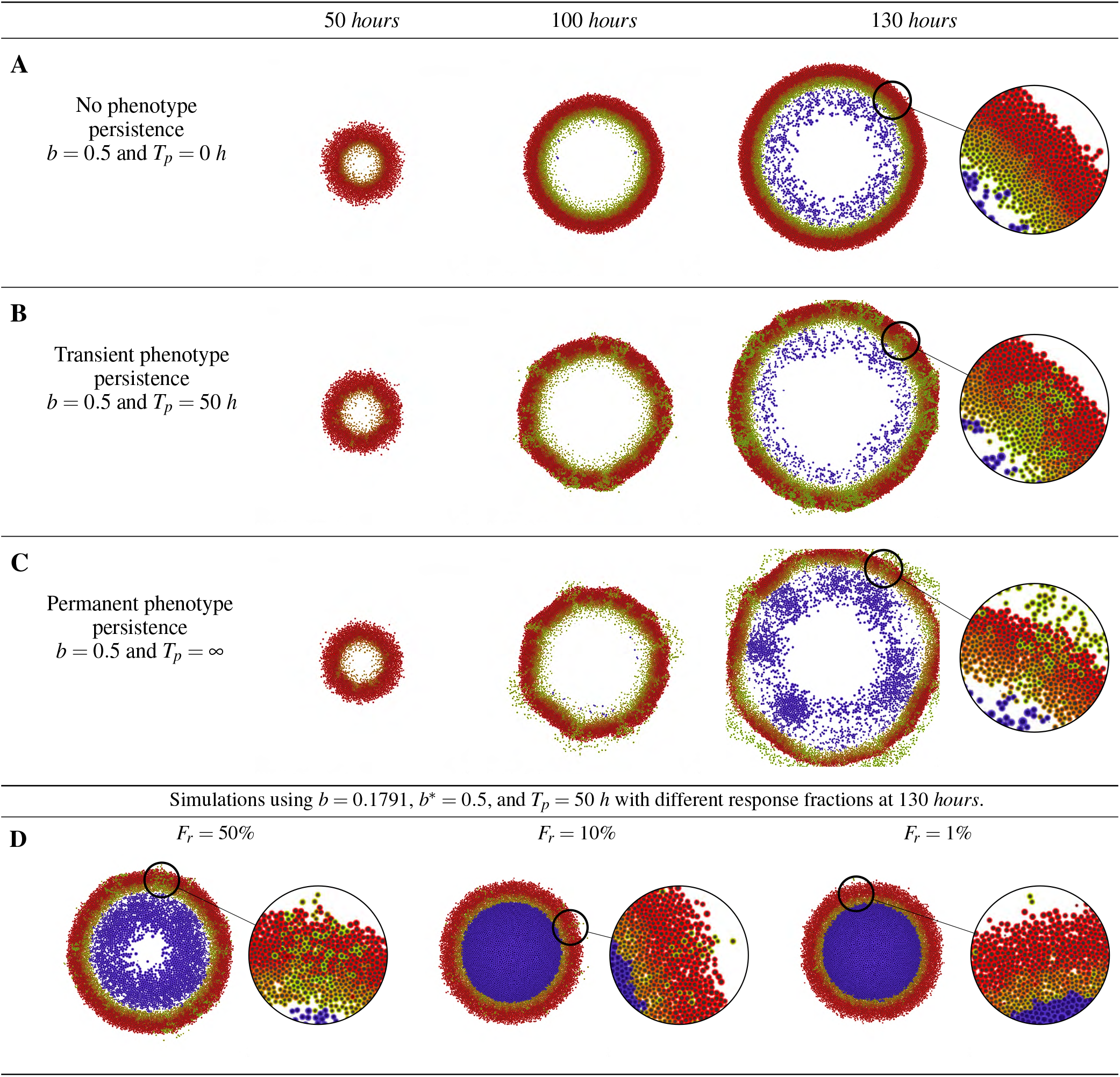
Impact of phenotypic persistence for directed migration. (A) Increasing migrational bias (*b*\* =0.5) without phenotypic persistence (*T_p_* = 0) drives invasion of GFP+ cells outside the tumor. (B) Under intermediate phenotypic persistence (*T_p_* = 50 *h*), GFP+ invasive “plumes” form inside the normoxic DsRed+ zones. Some GFP+ cells escape to invade the surrounding tissue. (C) Escape of GFP+ cells increases under permanent phenotypic persistence (*T_p_* = ∞). (D) For transient phenotypic persistence (*T_p_* = 50 h) and intermediate migrational bias (*b*\* = 0.5), decreasing the number of “responder” cells to 50%, 10%, and 1% gradually inhibits the formation of invasive plumes, although GFP+ cells continue to escape the tumor.

Because our experimental observations showed significantly higher displacement for a small fraction of GFP+ cells in the trajectory analysis, we hypothesized that invasion may be driven by a smaller subpopulation of “responder” cells: GFP+ cells that are capable of responding to hypoxia with a strong migratory phenotype. To computationally assess the potential impact of a responder subpopulation, we assumed that only a fraction of the GFP+ cells (*F_r_*) had a more directed motility along the O_2_ gradient with an elevated directional bias *b*\* = 0.5, while all other GFP+ cells retained the lower bias of *b* = 0.1791 obtained from our earlier calibration work. For simplicity, all GFP+ cells shared the same value of phenotypic persistence time (*T_p_* =50 *h*).

We found that as *F_r_* decreased from 100% to 50%, 10%, and 1%, the formation of invasive GFP+ plumes was suppressed, while the extent of necrosis increased. Without directed migration, the nonresponder GFP+ cells failed to escape low-oxygen regions, leading to increased necrotic death (Figure 4D). We note that even when only 1% of GFP+ cells could respond with increased migrational bias, GFP+ cells escaped the tumor to invade the surrounding tissue.

To better automate our analysis of the qualitative behavioral responses across the (*b*, *T_p_*, *F_r_*) parameter space, we developed an image processing pipeline to extract three morphometric features. For each simulation, we assessed (1) whether GFP+“plumes” were present, (2) whether GFP+ cells escaped the main tumor bulk, and (3) whether a necrotic core was present. See the supplementary materials for further detail. In our analysis, we found that these Boolean features were robust to stochastic variations in the simulations (e.g., from choosing a different random seed value). See the supplementary materials for an example where we show these features for several stochastic replica on a parameter set.

This automated analysis further confirmed that increasing phenotypic persistence time (*T_p_*) and motility bias (*b*\*) are associated with the formation of invasive plume structures and metastatic cell escape from the tumor bulk. Moreover, in scenarios with a substantial bias (*b*\* > 0.5), decreasing the fraction of responders (*F_r_*) is associated with increased necrosis and smaller tumor radii (Supplementary Figures 6,7,8). On the other hand, in scenarios of null bias (*b*\* < 0.1), a cell would have a low probability of leaving the hypoxic region, preventing the formation of post-hypoxic plumes (Supplementary Figures 7,8).

### Computationally predicted hypoxic patterns are observed *in vivo*

We performed parameter space exploration using the simulation model system to assess the emergent tumor behaviors under a variety of hypotheses on migration, phenotypic persistence, and heterogeneity in hypoxic response. For several conditions with phenotypic persistence (*T_p_* > 0), increased migrational bias (increased *b*\* for GFP+ cells), and heterogeneous hypoxic responses (*F_r_* < 100%), the model predicted the emergence of GFP+ plumes into DsRed+ well-oxygenated regions, and the formation of substantial necrotic cores. For instance, see in Figure 4, adopting *T_p_* = 50 *h*, *b*\* =0.5 and *F_r_* = 50%. These computationally predicted features are frequently observed in our *in vivo* tissue sections, where a necrotic core and GFP+ plumes were visible.

To further determine how well the computational model recapitulated the tumor biology, we visually compared a 2D tumor tissue section to our model output (Figure 5A). Strikingly, our model accurately predicted GFP+ invasive structures (Figure 5B, 5C). More specifically, we observed well defined persistent GFP+ plumes (1), as well as a second type of plume (2) that is larger and contains a mix of GFP+ and DsRed+ cells. Furthermore, the computational model also predicted DsRed+/GFP+ regions adjacent to the necrotic core (3) that were also observed experimentally. In both the simulation and experimental models, these double-positive cells begin to express GFP but the DsRed protein is not fully degraded indicating that these cells are currently hypoxic. All together, the data suggest that our computational model adopting *T_p_* = 50 *h*, *b*\* = 0.5 and *F_r_* = 50% most faithfully recapitulates the spatial localization of GFP+ cells *in vivo* during tumorgenesis, highlighting the formation of persistent invasive plumes.

**Figure 5.**
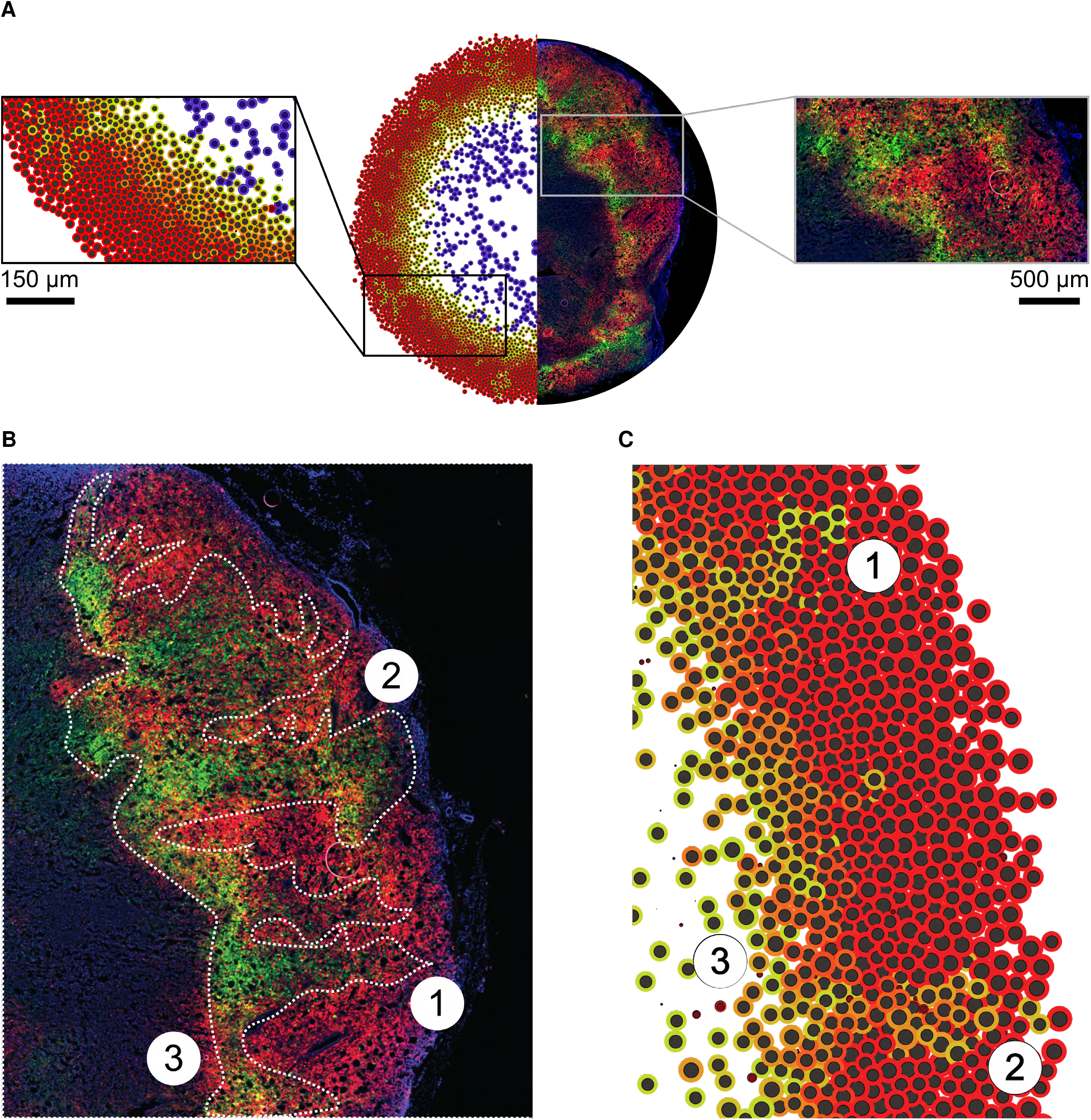
Computational model predicts the formation of post-hypoxic plumes. (A) Comparison of computational output to 2D tumor section. (B) Zoom of orthotopic tumor section. Dashed white outline highlights post-hypoxic plumes. (C) Zoom of computational output using *T_p_*= 50 h, *b*\* = 0.5 and *F_r_* = 50%. Numbers show analogous structures in both scenarios.

### Invasive plumes are exclusively driven by post-hypoxic cells

The GFP+ plumes appeared to extend into tumor regions with higher O_2_ content. Thus, we hypothesized that the GFP+ cells localized within the invasive plumes were no longer experiencing hypoxia, but rather were cells that had been previously hypoxic and were now reoxygenated. To experimentally test this, we utilized Hypoxyprobe, a well established antibody for the detection of hypoxic regions in tissue. Dual imaging of Hypoxyprobe and GFP expression allowed us to differentiate hypoxic cells (Hypoxyprobe+) from post-hypoxic cells (GFP+/Hypoxyprobe-) *in vivo*. While post-hypoxic cells (GFP+/Hypoxyprobe-) migrated into DsRed+ regions generating plumes, hypoxic cells (Hypoxyprobe+) localized adjacent to the necrotic core (Figure 6A). To visualize the hypoxic and post-hypoxic phenotypes in the computational model, we labeled the simulated cells localized alongside the perinecrotic region as Hypoxyprobe+ (Figure 6B). Remarkably, the computational model accurately recapitulated the observed biological differences. Together, these data suggest that the experimentally observed GFP+ invasive structures are composed exclusively of post-hypoxic cells (GFP+/Hypoxyprobe-), thus we termed them invasive post-hypoxic plumes.

**Figure 6.**
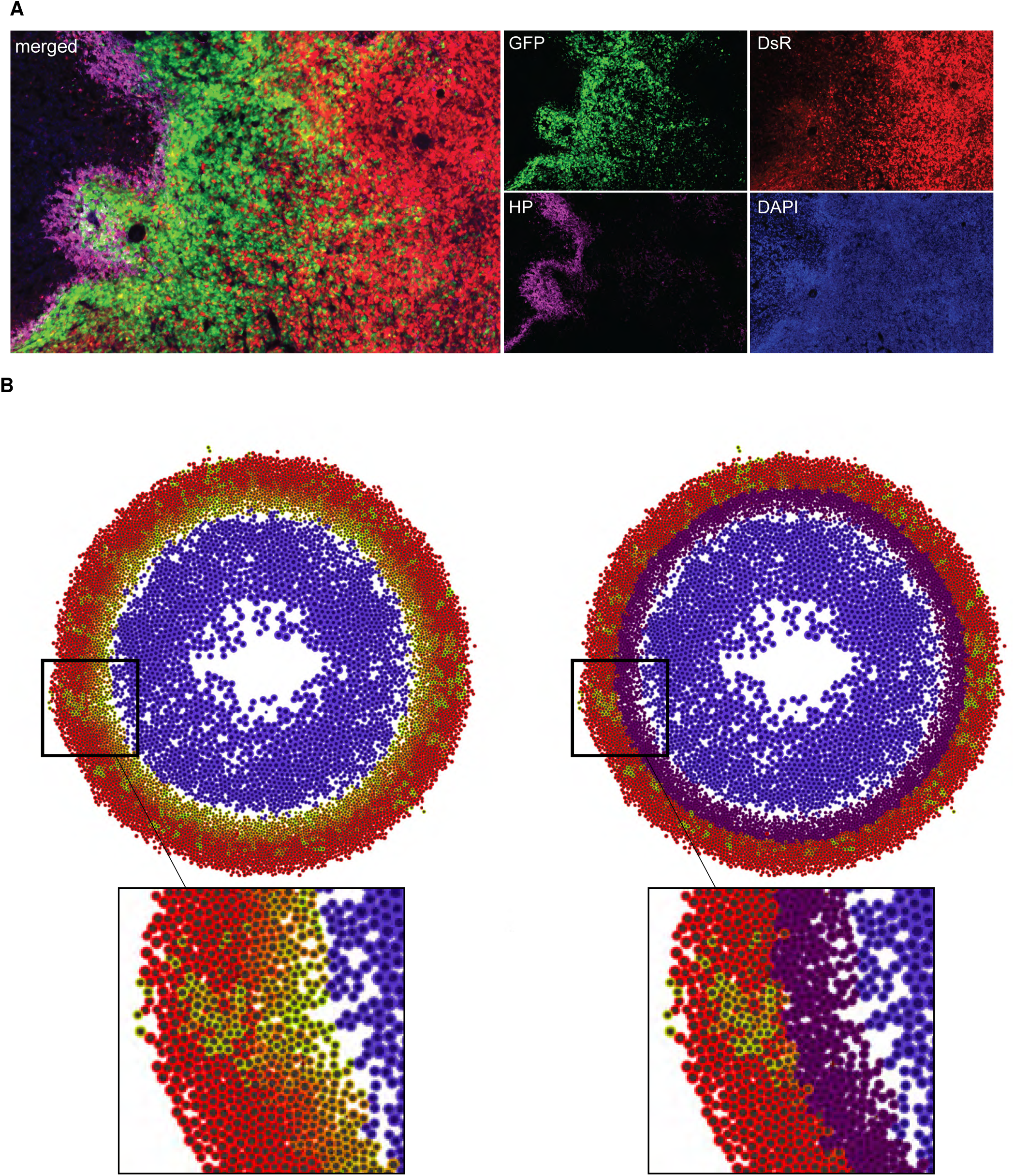
Comparison of Hypoxyprobe staining with hypoxia fate-mapping system. (A) Fluorescent image of orthotopic tumor section stained with Hypoxyprobe (HP). (B) Computational output without (left) and with (right) Hypoxyprobe (purple) marker.

### Computational model recapitulates intratumoral hypoxic patterns in 3D

We further tested whether the computational model’s predictions would extend to 3D by comparing simulations to a 3-D volume of tumor tissue. A 3D tumor (approximately 1,100 × 800 ×500 *μm*^3^) was reconstructed after CUBIC-tissue clearing, where the native fluorescence was imaged using confocal microscopy in a z-stack. The surface rendering of the 3D-reconstructed image demonstrated the presence of post-hypoxic plumes (Figure 7A). However, initial 3-D simulations (using the same parameter values as in 2D: *T_p_* = 50 *h*, *b*\* = 0.5, and *F_r_* = 50%) predicted less well-defined post-hypoxic GFP+ plumes than we observed experimentally (Figure 7B). We hypothesized that this apparent discrepancy arose due to differences between the computational and experimental geometries: the 3-D computational spheroid had a relatively small radius and hence high surface curvature, which contributed to dispersal of migrating GFP+ tumor cells. Under this hypothesis, 3-D simulations would better match experimental observations and earlier 2-D computational predictions for larger tumors in regions with flatter surface curvature.

**Figure 7.**
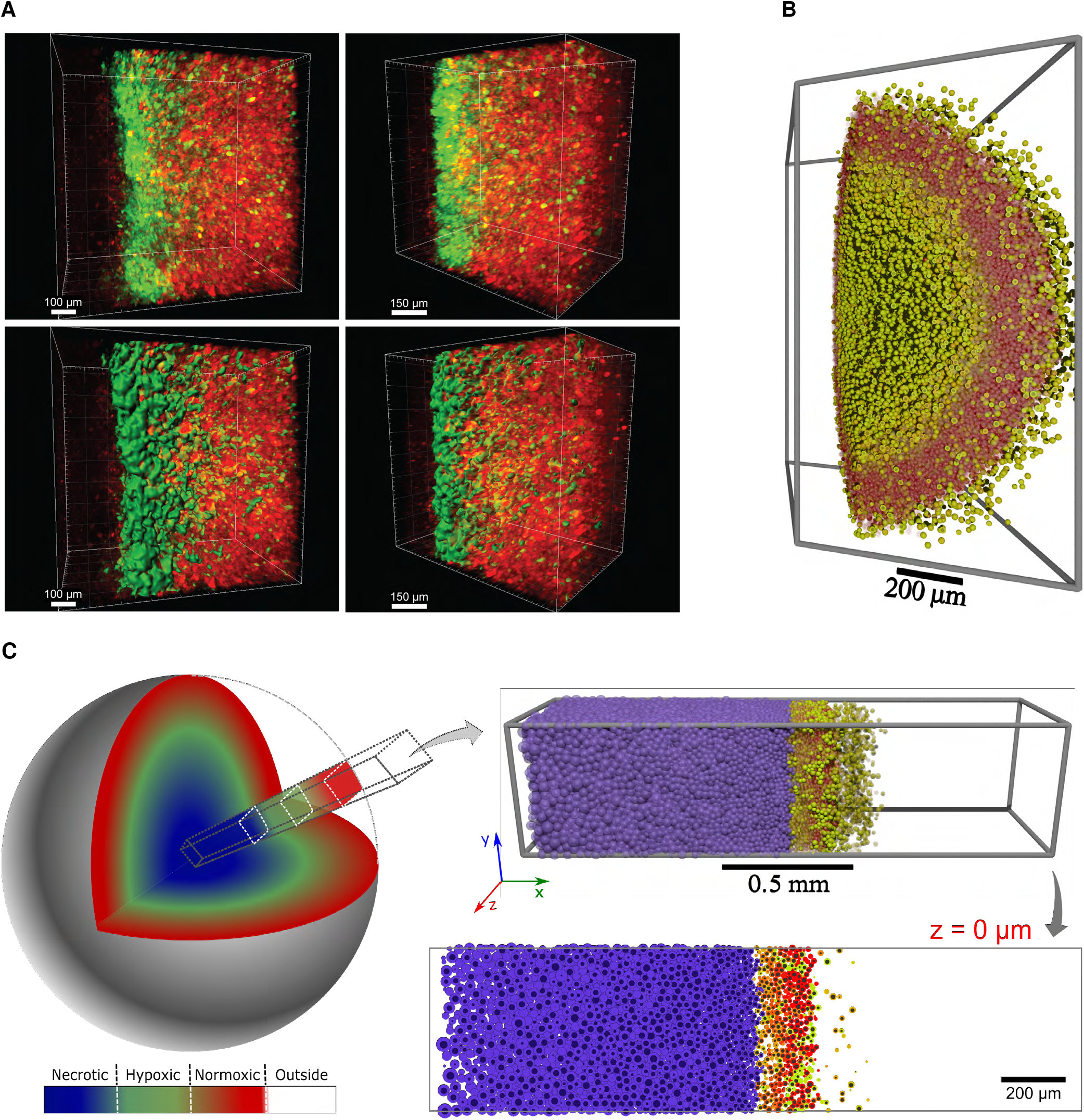
Computational model recapitulates intratumoral hypoxic patterns in 3D. (A) Reconstruction of cleared tumor section in the perinecrotic region. Raw reconstruction (top) and surface rendering of GFP+ signal (bottom). (B) Small spheroid generated by the 3D computational model. (C) Representation of a 3D approach *in silico* of a large tumor and low curvature (left) with highlight of a tumor radial projection in 3D and 2D (right).

We tested this by simulating a radial section of a larger 3-D tumor with flatter surface curvature: we modified the simulation domain to a 0.5 mm^3^ “core” (2 *mm* × 0.5 *mm* × 0.5mm) with no cell or substrate flux in the non-radial (*y* and *z*) directions, and with free cell and substrate flow in the radial (*x*) direction. (See this geometry in Figure 7C.) As hypothesized, necrotic regions and post-hypoxic GFP+ plumes were better defined in this 3-D geometry, and 2-D cross-sections were consistent with earlier 2-D simulation results. See Figure 7C.)

Taken together, the computational model accurately predicted the localization of cells exposed to hypoxia within a tumor and anticipated the formation of post-hypoxic plumes both in 2D and 3D. According to our model using *T_p_* = 50 *h*, *b*\* = 0.5 and *F_r_* = 50%, the resulting phenotype depends upon a balance of phenotypic persistence time, the fraction of responders, and migrational bias of the cellular movement, supporting the hypothesis of an enhanced migratory hypoxic/post-hypoxic phenotype.

## DISCUSSION

We combined a novel approach to fate-map cells that experience intratumoral hypoxia with advanced imaging and mathematical modeling to predict the dynamics of hypoxic and post-hypoxic cells during tumor progression. Investigating GFP+ (hypoxic/post-hypoxic) and DsRed+ (normoxic) cells in tumor tissue sections revealed that both cell populations have similar proliferation rates. On the other hand, GFP+ cells are more migratory *ex vivo* as compared to DsRed+ cells. Using these characteristics and the O_2_ distribution in a tumor, we modeled the dynamics of GFP+ cells with three key parameters: phenotypic persistence time (*T_p_*), fraction of responders (*F_r_*) and directional bias of the migrating GFP+ responders (*b*\*).

Our computational studies best recapitulated *in vivo* biological observations of (1) localization of GFP+ and GFP/DsRed double-positive cells to the perinecrotic regions of the tumor, (2) migration towards more oxygenated regions, (3) invasive GFP+ post-hypoxic plumes, (4) substantial necrosis, and (5) escape of GFP+ cells from the tumor bulk, when we defined phenotypic persistence (*T_p_* = 50 *h*), increased directional bias of migrating GFP+ cells (*b*\* = 0.5), and included heterogeneity in response to hypoxic stimuli (*F_r_* = 50%).

Furthermore, by examining the Hypoxyprobe marker for currently hypoxic cells both *in vivo* and *in silico* models, we determined that the GFP+ invasive structures were exclusively driven by post-hypoxic cells. The results support the conclusion that post-hypoxic plumes form due to a persistent migratory phenotype that develops when a cell becomes hypoxic and persists even when the cell is reoxygenated.

The *ex vivo* spheroid model used to determine cell motility consisted of measuring the displacement of tumor-derived cells (DsRed+ and GFP+) at the invasive front of spheroids embedded in a 3D collagen matrix. We observed enhanced speed, total displacement, and persistent time of GFP+ cells, suggesting that both migration and invasion contribute to cell motility in this model. In these experiments, we also observed that a fraction of GFP+ cells showed significantly higher displacement when compared to the remaining GFP+ cells, suggesting that only a subset of GFP+ cells acquire an increase in migration. To account for this in the model, we varied the directional bias for cell migration (*b*\*) for this responder subpopulation, and observed that higher values (*b*\* ~ 0.5) promoted the formation of post-hypoxic plumes. This phenomenon was surprisingly robust to perturbation: post-hypoxic plumes were consistently observed even when only 50% of the cells were responders (*F_r_* = 50%). However, experimentally-observed necrotic materials could only accumulate in the computational model when not all GFP+ cells were responders (*F_r_* < 100%). There are several potential biological explanations for this finding. First, the spatial localization of a cell within the O_2_ gradient in the tumor may cause heterogeneity in the level of migratory response. For example, cells localized closer to the necrotic core in a tumor experience lower O_2_ concentrations (<1%) and have higher levels of HIF expression, and therefore higher levels of HIF-dependent pathways that promote cell migration and invasion, such as RhoA-ROCK (Gilkes et al.,2014), ITGA5 (Ju et al., 2017), Notch (Chen et al., 2010), and miR-219-SMC4 (Chen et al., 2019), among others. Additionally, the observed invasion pattern suggests that cells localized in the invasive front must overcome matrix resistance by opening tracks, which would facilitate the invasion/migration of cells that follow, increasing their bias in a leader-follower-like manner (Haeger et al., 2015; Khalil and Friedl, 2010; Mayor and Etienne-Manneville,2016).

We explored a range of the phenotypic persistence time(s) and found that any sufficiently high value (50 hours or more in our studies) would result in the formation of GFP+plumes. This suggests that the formation of post-hypoxic plumes is associated with a persistent phenotype that is maintained in GFP+ tumor cells for a considerably long duration. Although HIF-1 protein levels return to baseline within minutes after reoxygenation to room air (≈20% O_2_) (Jaakkola et al., 2001; Semenza and Wang, 1992), the time required for the protein products of HIF-regulated genes to return to baseline will depend upon the half-life of the protein. Moreover, the hypoxic cells within a tumor encounter a slower reoxygenation as they migrate towards the oxygenated areas, and experience a lower O_2_ gradient (1% to 6% O_2_) causing prolonged HIF-signaling when compared to *in vitro* studies. Therefore, post-hypoxic cells within *T_p_* = 50 *h* will retain an increased level of HIF-target gene and protein expression which could, in part, explain the lengthy persistence time. Indeed, we previously demonstrated that cells exposed to intratumoral hypoxia, but not short-term *in vitro* hypoxia, retain the increased expression of a subset of hypoxia-regulated genes, even upon reoxygenation and long-term culture (Godet et al., 2019). Further studies would be necessary to establish whether the expression of one or multiple genes plays a causal role in phenotypic persistence. However, it is tempting to speculate that the ‘hypoxia memory’ genes LOX or DNAH11 could be implicated, as they can regulate cell movement (Lai et al., 2016; Payne et al., 2005).

Interestingly, we noticed that the observed computational plumes seem narrower than the biological ones. While our *in vivo* tumors reach diameters of 10 mm, our computational model reached a radius of about 0.6 mm. Therefore, it could be expected that larger tumors would give origin to larger plumes. The computational time to model a tumor with a diameter of 10 mm is not feasible. Moreover, imposing dynamics of leader-follower / collective invasion that can potentially be happening *in vivo* could drive larger computational plumes. Lastly, the *in vivo* time scale is larger than the computational one, and therefore, proliferation of GFP+ cells might be contributing to larger plumes as well. Overall, our computational model recapitulates GFP+ plume formation, and perhaps larger tumors and longer time-scales could contribute to generate larger plumes.

It is important to note that the MDA-MB-231 orthotopic tumors modeled in this work are uniformly radial and homogeneous. To test if our observation of GFP+ plume formation would still occur in heterogeneous environments, we performed a simulation varying the location and number of O_2_ sources. In order to avoid non-physical behaviour, we imposed restrictions on cell proliferation and migration in regions of higher cell density (See Supplementary materials for details). In this heterogeneous scenario, GFP+ plumes were still observed, mainly around more oxygenated regions (Supplementary Fig. 9) This suggests that our observation of GFP+ plume formation is robust and O_2_-dependent and not simply a consequence of radial growth. Human breast tumors have a slower growth rate, and therefore have additional time for angiogenesis to occur. Angiogenesis results in a random distribution of functional vasculature within the tumor that can cause heterogeneous O_2_ gradients (Vaupel et al., 2007). Together, these factors promote multiple microregional O_2_ gradients that exist within a larger O_2_ radial gradient across the tumor. It would be interesting to incorporate an intermittent subscale O_2_ distribution, by modeling angiogenesis and blood supply dynamics. Transient changes in O_2_ gradients that occur locally have been reported but occur in the range of 1.3-2.6%(Matsumoto et al., 2010). However, the overall O_2_ distribution within a viable rim of 2 cm in a large tumor ranges from 0 to 6% O_2_ (Godet et al., 2019) suggesting that the radial O_2_ gradient observed in our experimental tumors is the largest contributor to plume formation. Therefore, we hypothesize that the model would reveal similar radial O_2_ gradients with GFP+ plumes formation on both macro and microscales.

In the future, we plan to extend the mathematical model to include cell signaling and to more accurately match the dynamics such as O_2_ distribution, proliferation and tissue and cell mechanics. The implementation of new calibration protocols to leverage High Performance Computing (HPC) and ABC techniques will be crucial to simultaneously evaluate all these parameters. Ultimately, we will also aim to advance our single-cell tracking and plan to include additional cell cycle markers. The labeling and characterization of blood vessels in tumor sections (CD31 or dextran) could be used to assess angiogenesis. In addition, the contribution of the extracellular matrix and the leader-follower dynamics will be important to consider. The HIF-response could be incorporated in our computational model by building a network of protein interactions and considering their half-life times, which could further develop our studies in the hypoxic/post-hypoxic phenotypes.

In summary, we developed a computational model based on the characterization of normoxic, hypoxic and post-hypoxic cancer cell phenotypes that accurately predicted the formation of novel structures that we termed ‘invasive post-hypoxic plumes’. Although it is well established that hypoxia promotes migration and invasion *in vitro* during exposure to hypoxia, we show that post-hypoxic cells retain a migratory phenotype that contributes to plume formation. While at the macroscopic level the invasive plumes *appear* to be driven by collective invasion (a coordinated and guided migration) (Yang et al.,2019), our computational findings demonstrate that these structures can spontaneously arise without any direct cell-cell coordination. The results show the potential for combining novel experimental approaches with computational modeling: the hypoxia fate mapping system rendered the previously unobserved post-hypoxic plumes visible, computational modeling allowed us to probe the underlying dynamics to suggest hypotheses that can generate the structures, and targeted experimental analyses were able to confirm the computational predictions.

## STAR★METHODS

### Experimental Methods

#### Cell culture

Mycoplasma-free human breast cancer cell line MDA-MB-231 was obtained from the American Type Culture Collection (ATCC) and maintained in DMEM (Sigma-Aldrich) with 10% FBS (Corning) and 1% penicillin/streptomycin (Invitrogen). An InvivO_2_ workstation (Baker) with an ICONIC (Baker) electronically controlled gas-mixing system was used to achieve hypoxic conditions in a controlled environment at 37°C and 75% humidity, and balanced at 0.5% O_2_, 5% CO_2_, and 94.5% N_2_.

#### Hypoxia fate-mapping system

The design of the hypoxia fate-mapping system has been described previously (Godet et al., 2019). Briefly, the loxp-DsRed-loxp-eGFP sequence was PCR amplified from the plasmid pMSCV-loxp-DsRed-loxp-eGFP-Puro-WPRE (#32702, Addgene). This sequence was then ligated into the pENTRA1 vector and the Gateway System TM (Invitrogen) was used to recombine the pENTR1A shuttle vector with pLenti CMV/TO Zeo DEST (644-1) (#17294, Addgene) generating lentiviral vector 1. The sequence GTGTACGTG (1HRE) spaced with random 5 base pairs of nucleotides was used to generate tandem copies of 4 HREs that were directly synthesized (IDT) as gBlocks (Coralville). The CRE-ODD nucleotide sequence was also independently synthesized as a gBlock (IDT). The pRRLSIN.cPPT.PGK-GFP.WPRE (#12252, Addgene) was digested to remove the PGK promoter and EGFP cassette. In-Fusion cloning (Clonetech) was used to ligate the 4-HRE gBlock, the *β*,-globin gBlock and the CRE-ODD gBlock into the linearized pRRLSIN.cPPT.PGK-GFP.WPRE vector to generate vector 2. Finalized lentiviral vector 1 encoding CMV-loxp-DsRed-loxp-eGFP or lentiviral vector 2 encoding 4xHRE-MinTK-CRE-ODD were co-transfected with plasmid psPAX2 (#12260, Addgene) and plasmid pMD2.G (#12259, Addgene) into 293T cells using Polyjet (SL10088, Signagen). Filtered viral supernatant generated from lentiviral vector 1 was collected 48 h post-transfection and added to MDA-MB-231 cells with 8 μg/mL polybrene (Sigma–Aldrich) overnight. After 24 h in fresh media, zeocin (Invitrogen) was added to the medium of cells for selection (100 μg/mL). Following selection, cells were transduced with lentivirus from lentiviral vector 2 encoding 4xHRE-MinTK-CRE-ODD. The cell lines were sorted, single-cell cloned and screened by image analysis and flow cytometry.

#### Animal model

Female 5- to 7-week-old NOD-SCID Gamma (NSG) mice were used according to protocols approved by the Johns Hopkins University Animal Care and Use Committee. Mice were anesthetized by the intraperitoneal injection (i.p.) of 100 mg/kg Ketamine, 16 mg/kg Xylazine, Vet One. 2×10^6^ MDA-MB-231 hypoxia fate-mapping cells were injected into the mammary fat pad (MFP) closest to the second nipple. Mice were i.p. injected with 1.25 mg of pimonidazole in saline (12.5 mg/ml) (Hypoxyprobe^™^-1) 1 h prior to sacrificing. Tumors were excised at various time points, formalin fixed (Sigma-Aldrich) for 1 h, saturated in 30% sucrose (Sigma-Aldrich) at 4°C overnight, embedded in OCT media (Fisher Scientific), frozen in liquid nitrogen, sectioned via a cryotome CM1100 (Leica), and mounted onto Superfrost Plus Microscope Slides (Fisher Scientific). Tumor tissue sections were stained with DAPI (1:1000 for 15 min, RT) and mounted with anti-fade solution. To assess the entire cross-section of the tumor, slides were imaged with an Olympus (UPLFLN 4X) objective using Cytation 5 microscope (BioTek Instruments). Multiple image tiles were linearly stitched with Gen5 Software (BioTek Instruments).

#### Oxygen measurements

*In vivo* O_2_ measurements were conducted on mouse tumors using a fixed-needle (0.9 mm / 230 *μm*) REDFLASH mini-sensor (FireStingO_2_, Pyroscience) mounted on a manual micrometer. The probe was inserted approximately 2 mm into the tumor, and after recording the hypoxic level, it was slowly retracted and O_2_ measurements were recorded at 0.5 mm intervals until reaching the tumor edge. The needle-probe was carefully washed and calibrated to atmospheric O_2_ before each measurement.

#### Immunofluorescence staining

Tumor tissue sections were washed 3x with PBS-T (PBS, 0.1% Tween) and briefly air-dried. Using a hydrophobic pen (Dako, Angilent), a circle was drawn around the tumor sections, and they were then incubated with 1% Triton-X at room temperature for 10 min in the dark. After a new wash with PBS-T, the tissue sections were incubated with 2% BSA at room temperature for 30 min in the dark. The Alexa Fluor 647 Mouse anti-Ki-67 antibody (BD Biosciences, #558615) was used at 1:50 dilution in BSA at room temperature for 1.5 h in the dark. New PBS-T wash was followed by staining with DAPI at 1:1000 dilution in PBS at room temperature for 15 min in the dark. Slides were washed and mounted with anti-fade solution. Images were taken with an Olympus (UPLFLN 10X) objective using Cytation 5 microscope (BioTek Instruments). Image analysis was performed by determining the ratio of Ki67+ nuclei over total nuclei in each RFP and GFP channels using the image calculator tool available on ImageJ.

#### Spheroid model

Cell spheroids were formed in round-bottom 96-well tissue culture plates as described previously (Valencia et al., 2015), (Ju et al., 2017). Briefly, a mixture of (1×10^4^) cells, 80-90% unlabeled cells and 10-20% hypoxia fate-mapping cells derived from orthotopic tumors, were plated per well in spheroid formation media 1:1 DMEM (Sigma-Aldrich) and Methocult H4100 (STEMCELL Technologies). Plates were centrifuged at 1200 rpm for 7 min, twice. After 72 h incubation, each spheroid was transferred to a petri dish, where they were individually isolated with the collagen solution (2 *mg*/*ml*) and quickly transferred to the center of a semi-crosslinked collagen gel in a 96 well-plate at 37°C. After complete crosslinking, warm media was added. Following 4 days in culture, spheroids were imaged in an environmentally controlled microscope every 15 min for 16 h using an Olympus (UPLFLN 4X) objective in Cytation 5 (BioTek Instruments). Cell trajectories were tracked using MetaMorph software to obtain x,y coordinates at each time.

#### CUBIC tissue clearing

Tumor tissue clearing was achieved by following the CUBIC protocol (Susaki et al., 2015), and CUBIC-I (water 35% wt, urea 25% wt, Quadrol 25% wt, Triton-X 15% wt) and CUBIC-II (sucrose 50%, urea 25%, water 15%, triethanolamine 10%) solutions were made prior to use. Tumors resected at day 20 were fixed in formalin at 4°C overnight. Tumors were then incubated with 1:1 CUBIC-I/water overnight at 37°C and 60 RPM in an orbital shaker. On the following day, tumors were transferred to CUBIC-I and kept at 37°C and 60 RPM in an orbital shaker until desired clearness was achieved (approximately 2.5 weeks), with CUBIC-I being replaced every other day. Once clearing was satisfactory, tumors were transferred to CUBIC-II for 2 days. Tumors were imaged in mineral oil using confocal microscopy to obtain z-projections of tumor sections with approximately 500 *μ*m depth using a 10X/0.45 PlanApo (dry, no DIC) objective in a Zeiss LSM780-FCS microscope. Z-stacks spaced at 6.3 *μ*m intervals were processed into a 3D image via Imaris version 9.2 (Bitplane) and 3D surface rendering was used to visualize the GFP+ invasive structures in 3D.

### Computational Methods

#### PhysiCell: agent-based cell modeling

The computational model was developed using PhysiCell, an open-source C++ framework allowing the construction of multicellular models at various scales in 2D and 3D (Ghaffarizadeh et al., 2018) on a broad variety of computing platforms. In this structure, each cell has its phenotypic and physical properties individually characterized through an agent-based model. Generally, cell dynamics are partially determined by diffusion of substrates such as oxygen, glucose, and growth factors in the microenvironment. To model the distribution of these substrates in the tissue environment, PhysiCell uses an open-source biological diffusion solver, BioFVM (Ghaffarizadeh et al., 2016). The coupling between these computational tools is defined by the consumption and secretion of substrates by each cell. PhysiCell has been applied to a broad variety of multicellular problems, such as oncolytic virus therapy, cancer immunology, tissue mechanics, infection dynamics and tissue damage, and drug screening, among others (Getz et al., 2020; Ghaffarizadeh et al., 2018; Jenner, 2019; Ozik et al., 2019; Risner et al., 2020; Wang et al., 2020). Please see Ghaffarizadeh et al. (2018) for further computational details and performance testing.

#### Computational model

Using our biological observations, we applied PhysiCell (version 1.6.1) to develop a mathematical model representative of tumor progression that incorporates the cell phenotype, location, and exposure to spatially dependent O_2_ concentrations. In this model, we differentiate the normoxic, hypoxic and necrotic state of the cancer cell based on the level of pO_2_. The O_2_ concentration is distributed in the environment using the standard transport equations from BioFVM and PhysiCell (Ghaffarizadeh et al., 2016, 2018). Here, we define the pO_2_ as σ. In regions outside the tumor, we set the O_2_ concentration to a (maximum) far-field value, denoted by 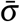.

A change in phenotype from normoxic to hypoxic occurs only in response to a decrease in O_2_ availability below the threshold, σ_*H*_. When the *O*_2_ level decreases further, below a *σ_T_* threshold, the cell is modeled necrotic (or dead). (Full details on the necrosis model can be found in Ghaffarizadeh et al.(2018).) When a hypoxic cell migrates to a region with a high supply of oxygen, we characterized it as a post-hypoxic or reoxygenated cell. The concentration of pO_2_ also contributes to the rate of cellular proliferation in this model. We use the built-in Ki67 basic model of the PhysiCell, where each cell has two states according to its current stage in the cell cycle. The first state [*Ki*67–] represents cells with absence of Ki67 expression, and the second state [*Ki*67+] represents positive expression of Ki67, in which cells have already started the proliferation process.

In our hypoxia fate-mapping system, all cells begin by expressing DsRed, and upon exposure to hypoxia they permanently switch from DsRed to GFP expression. The process of transitioning from DsRed to GFP protein expression is modeled through a system of two ordinary differential equations (ODEs), in which given a set of (normalized) genes **G** = (*G*_0_, *G*_1_) encoding the proteins, the (normalized) protein concentration in each cell is denoted by:

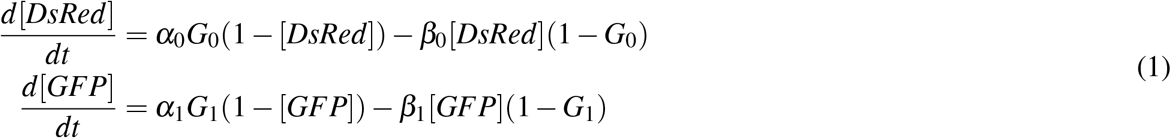

where *α_i_* is a protein production rate and *β_i_* is a protein degradation rate. (Note that in this normalized model, each protein *i* decays exponentially with rate *β_i_* when its gene expression is 0, and it tends towards a steady state value of 1 when its gene expression is 1.) We calibrated the production and degradation rates of GFP and DsRed, respectively, based on our experimental data. As an initial condition, we assumed that [*DsRed*] (0) =1 and [*GFP*] (0) = 0. As a simplifying assumption, we define **G** = (1,0) as gene expression of normoxic cells that never experienced hypoxia, and **G** = (0,1) as permanent gene expression of cells that have been subjected to hypoxia. It is important to note that GFP expression is permanent once cells experience hypoxia due to the gene editing mechanism (Figure 2A, 2B).

Cellular motility is characterized in the model by four intrinsic properties: persistence time (τ: the mean time that the cell maintains its current speed and direction); migration speed (*s*: the speed of cellular migration in the absence of other forces); the migration bias direction (**d**: the preferred direction of migration), and migration bias (*b*: a value between 0 and 1 that represents the extent that cell trajectory mimics Brownian motion). Zero bias (*b* = 0) represents purely Brownian motion, whereas *b* = 1 is completely deterministic motion along its preferred direction **d**. (See Ghaffarizadeh et al. (2018) for further details, or an online motility demonstrator at https://nanohub.org/tools/trmotility.) In the present model, we use the O_2_ gradient **∇**σ as the preferred direction of motility for biased migration (**d** = ∇σ).

Based on our biological observations (Supplementary figures 1b,c), we assume that DsRed+ and GFP+ cells have identical proliferation capacity under identical conditions. In contrast, cell migration has specific characteristics for each type of cell. As seen in the biological experiments, GFP+ cells have a more pronounced migratory phenotype.

#### Approximate Bayesian Computation

Model parameters were calibrated using an Approximate Bayesian Computation (ABC) method. ABC is a class of inference techniques based on Bayesian statistics without the explicit construction of the likelihood function, and this terminology has been used since 2002 (Beaumont et al., 2002). This approach has become quite popular and has been applied to problems of parametric inference and model selection. Particularly, this technique has been widely used in the analysis of complex problems in the biological sciences (such as systems biology, ecology, and others) (Sunnåker et al., 2013).

In a general Bayesian inference procedure, we want to estimate the parameters associated with the model, such that the observational values are matched with the model outputs. Here, we apply an ABC method proposed by Marjoram et al (Marjoram et al., 2003), named Likelihood-free Markov chain Monte Carlo (LF-MCMC). We compute an approximation of the posterior distribution of the model parameters, *π_LF_*(*θ* |*y_o_*), based on an a priori distribution of the parameters, *π*(*θ*), and observational data, *y_o_*. Thus, an approximate version of Bayes’s theorem (Sisson and Fan, 2011) is:

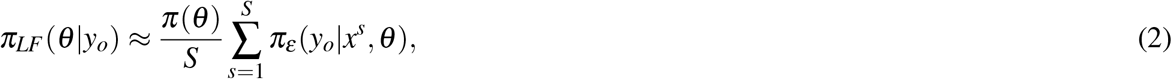

where *x*^1^, ⋯, *x^S^* are independent draws from the model *π*(*x*|*θ*) and *ε* is a tolerance value of the residual between the experimental data (*y_o_*) and the model output (*x*). It is important to note that when *ε* is small enough, using a conventional metric, the approach to posterior distribution is more accurate. However, choosing a small epsilon requires a high computational effort. In this work, for simplicity, we used the metric induced by the *ℓ*_2_ norm and estimate epsilon values based on the observational data.

#### nanoHUB platform

The computational model built is available for simulations on the nanoHUB platform (Madhavan et al., 2013). The nanoHUB platform is an open and free environment that offers several services aimed at the dissemination of science and engineering products. In particular, it offers a toolkit for the development of graphical user interfaces (GUIs) that allow greater accessibility and reproducibility of software.

#### Code availability

All code is available at: https://github.com/PhysiCell-Models/breastcancer-hypoxia.

We used *xml2jupyter* (Heiland et al., 2019) to generate a Jupyter-based graphical user interface (GUI) from our PhysiCell model. An online interactive version of the model is available at: https://nanohub.org/tools/pc4tumorhypoxia.

## ACKNOWLEDGMENTS

We would like to acknowledge Jisu Shin in the Gilkes lab for her work on cell tracking in the spheroid model. This collaborative work is part of the Applied Mathematics in Germinating Oncology Solutions (AMIGOS) workshop hosted by The Jayne Koskinas Ted Giovanis Foundation for Health and Policy and by the Breast Cancer Research Foundation.

Research in the Macklin lab is supported by U01-CA232137 (NCI), 1720625 (NSF), 1818187 (NSF), the Breast Cancer Research Foundation, and the Jayne Koskinas Ted Giovanis Foundation for Health and Policy.

Research in the Gilkes lab is supported by U54-CA210173 (NCI), Susan G. Komen Foundation (CCR17483484), Cindy Rosencrans Fund for Metastatic Triple-Negative Breast Cancer, The Emerson Collective, The Jayne Koskinas Ted Giovanis Foundation for Health and Policy and the SKCCC Core Grant (P50CA006973(NCI)).

This research was supported in part by Lilly Endowment, Inc., through its support for the Indiana University Pervasive Technology Institute.

## AUTHOR CONTRIBUTIONS

H.L.R. was responsible for code development and computational data assembly. I.G. was responsible for biological research design and execution; F.K., J.M., K.K. and S.B. assisted with code development. D.M.G. was responsible for biological research design and supervision. P.M. was responsible for computational model conception and supervision. H.L.R., I.G., D.M.G. and P.M. wrote and edited the manuscript.

