## Supplementary Materials for "A persistent invasive phenotype in post-hypoxic tumor cells is revealed by novel fate-mapping and computational modeling"

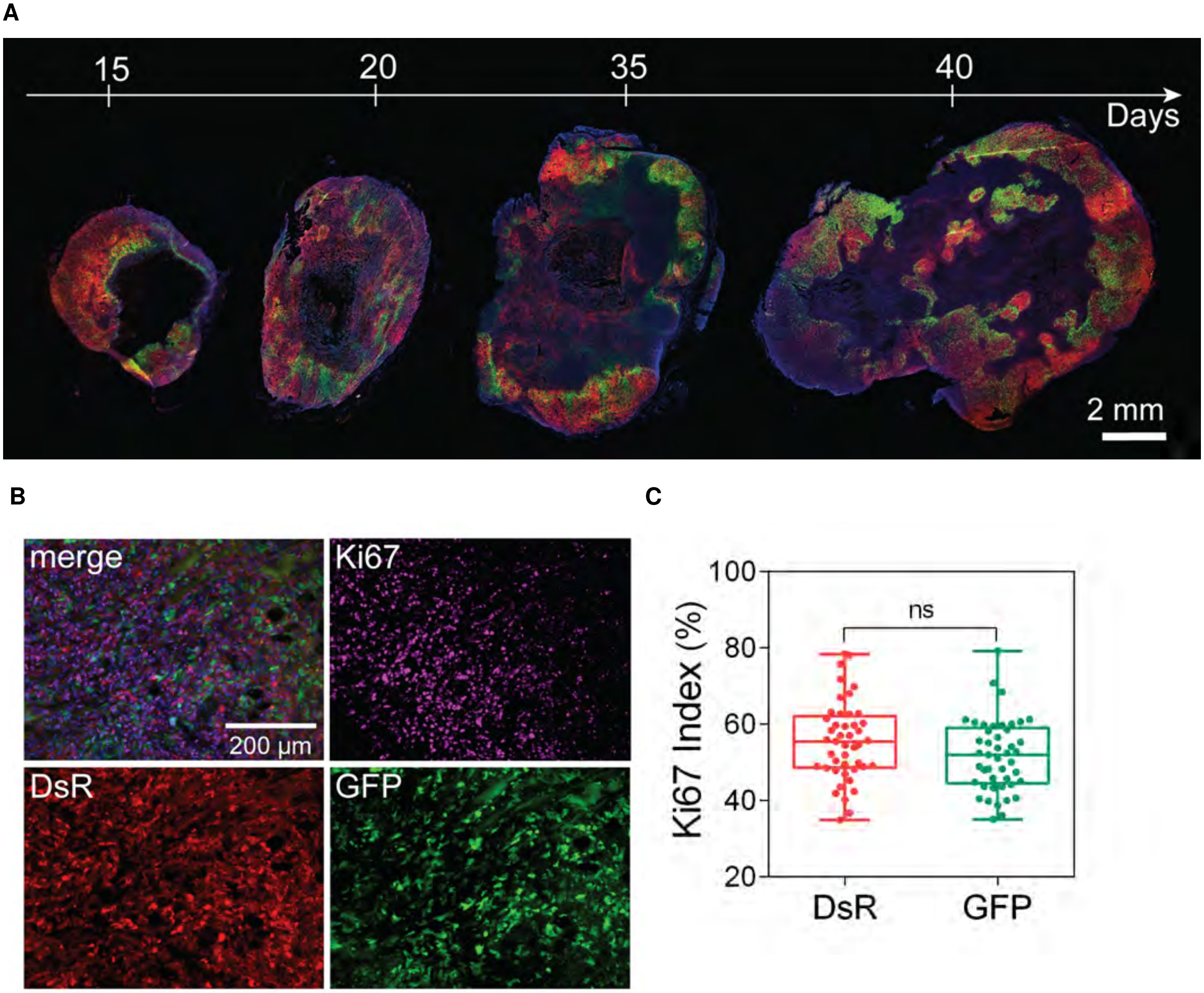


**Figure 1. Fate-mapping intratumoral hypoxia and proliferation analysis**. (A) Fluorescent images of the full cross-sections of orthotopic tumors derived from MDA-MB-231 hypoxia fate-mapping cells. Tumors were excised at days 15, 20, 35 and 40 of the time course. (B-C) Representative fluorescent image (B) and quantification (C) of Ki67 staining in DsRed+ and GFP+ cells in a tumor section.

| Parameter | Meaning | Value | Reference |
| --- | --- | --- | --- |
| $U$ | oxygen consumption rate by cells | $10\ min^{-1}$ | Ghaffarizadeh *et al.* 2016 |
| $D$ | oxygen diffusion coefficient | $10^5\ \mu m^2/min$ | Ghaffarizadeh *et al.* 2016 |
| $\lambda$ | oxygen natural decay | $0.1\ min^{-1}$ | Ghaffarizadeh *et al.* 2016 |
| $\sigma_0$ | initial oxygen pressure | $45.94\ mmHg$ | Estimated |
| $\bar{\sigma}$ | oxygen pressure at the edges of tumor | $45.94\ mmHg$ | Godet *et al.* 2019 |
| Initial_radius | radius of the initial tumor | $250\ \mu m$ | Estimated |
| $\bar{r}_{01}$ | trasition rate from Ki67- to Ki67+ | $3.63 \times 10^{-3}\ min^{-1}$ | Ghaffarizadeh *et al.* 2018 |
| $r_{10}$ | trasition rate from Ki67+ to Ki67- | $1.07 \times 10^{-3}\ min^{-1}$ | Ghaffarizadeh *et al.* 2018 |
| $\sigma_S$ | oxygen pressure threshold to signal proliferation saturation | $38\ mmHg$ | Estimated |
| $\sigma_T$ | min oxygen pressure threshold to signal proliferation | $6\ mmHg$ | Estimated |
| $\sigma_H$ | oxygen pressure threshold to signal hypoxia | $10\ mmHg$ | Godet *et al.* 2019 |
| $\alpha_i$ | protein production rate | $4.8 \times 10^{-4}\ min^{-1}$ | Estimated |
| $\beta_i$ | protein degradation rate | $6.9 \times 10^{-5}\ min^{-1}$ | Estimated |

**Table 1. Parameter values utilized in the simulations of the computational model.**

### Supplementary results

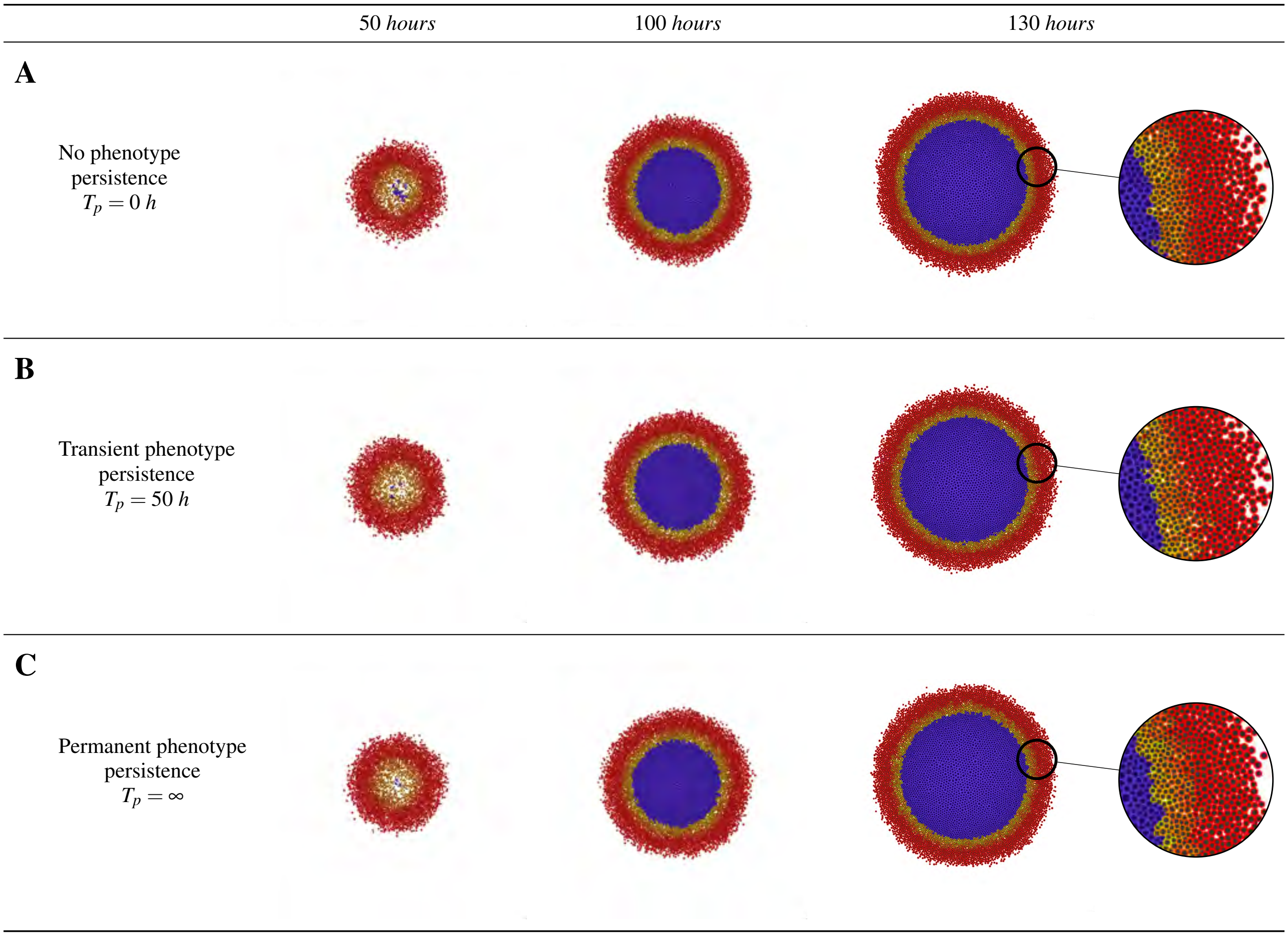


**Figure 2. Impact of phenotypic persistence time with migratory bias fixed ($\mathbf{b = 0.1791}$).** (A) No phenotypic persistence ($T_p = 0$). (B) Intermediate phenotypic persistence ($T_p = 50\ h$). (C) Permanent phenotypic persistence ($T_p = \infty$).

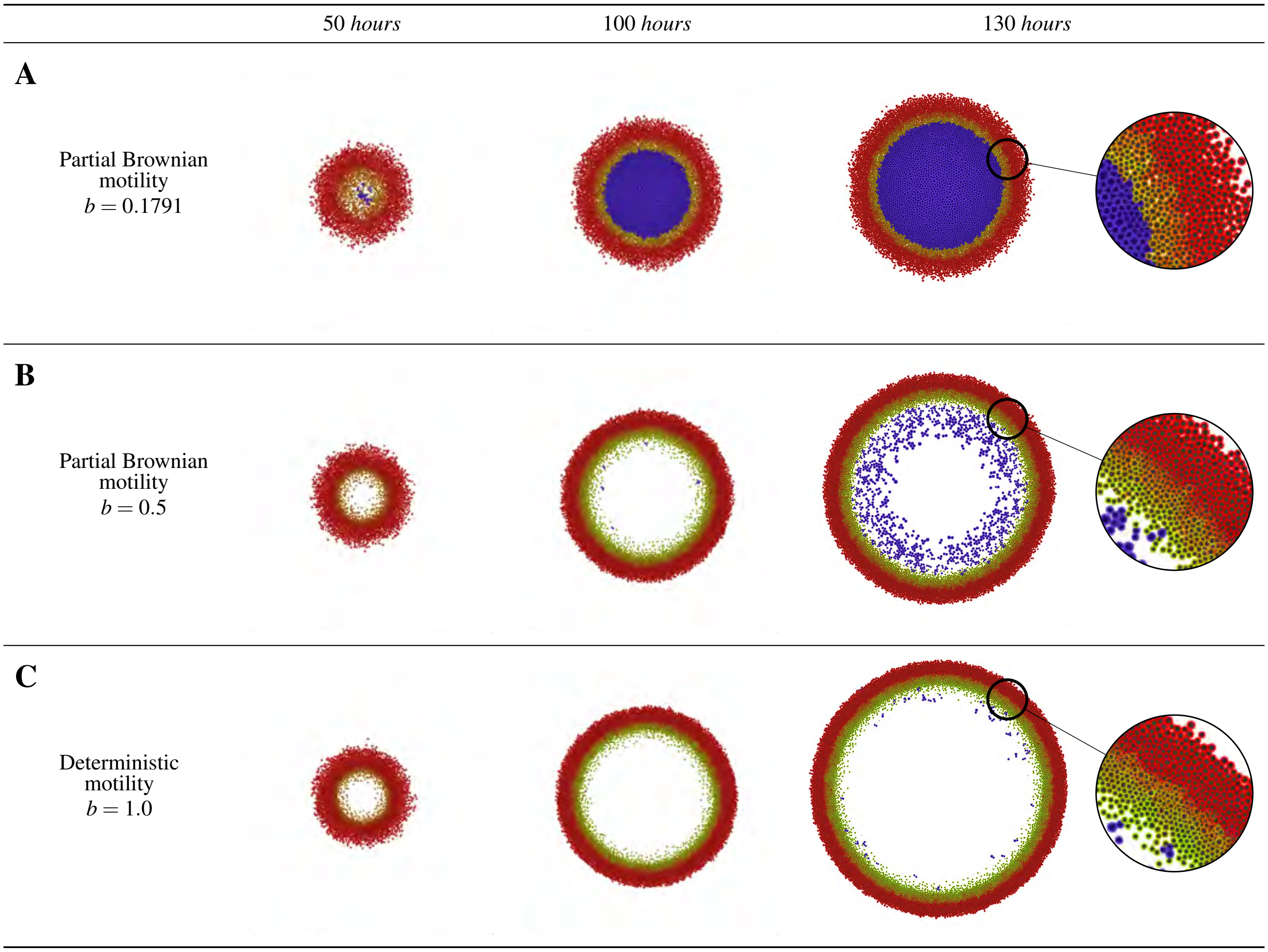


**Figure 3. Impact of migratory bias without phenotypic persistence ($\mathbf{T_p = 0}$).** (A) The motility bias generates a partial Brownian motility ($b = 0.1791$). (B) Intermediate motility bias achieves a more directed migration ($b = 0.5$). (C) Completely polarized movement ($b = 1.0$).

#### Image classifier

We built an image Boolean classifier to define the occurrence of plumes, escaping cells, and necrotic core (see the example in Figure 4). To check whether plumes formed, we extract the green cells from the image and then select bigger contour of the cell cluster. We fit that contour by an ellipse $\mathcal{E}_G$ and by a piecewise polynomial function $\mathcal{F}_G$. Based on the distance of the vertices of the function $\mathcal{F}_G$ and ellipse $\mathcal{E}_G$, we define whether there is a plume or not, according to tolerance $\varepsilon_P$ (Figure 4B). For the cell escape test, we fit an ellipse $\mathcal{E}_T$ to the tumor and search for green cells outside that ellipse, and then we verify if the sum of these cells' areas is greater than $\varepsilon_S$ (Figure 4C). The existence of the necrotic nucleus in the image occurs when the area fraction of necrotic cells is greater than $\varepsilon_N$ (Figure 4D). In general, the tests are defined as:

$$\text{Plumes:} \quad d(v,\mathcal{E}_G),\ v \text{ vertex of } \mathcal{F}_G \quad \begin{cases} True, & \text{if } d(v,\mathcal{E}_G) > \varepsilon_P \\ False, & \text{else} \end{cases}$$

$$\text{Cell escape:} \quad A = \sum A_i,\ A_i \text{ area of the GFP+ cell } i \text{ outside of } \mathcal{E}_T \quad \begin{cases} True, & \text{if } A > \varepsilon_S \\ False, & \text{else} \end{cases}$$

$$\text{Necrotic core:} \quad F_N = \frac{\text{Necrotic area}}{\text{total area}} \quad \begin{cases} True, & \text{if } F_N > \varepsilon_N \\ False, & \text{else} \end{cases}$$

in which $d$ is distance function. In this work, we used $\varepsilon_P = 30\ px$, $\varepsilon_S = 20\ px$, and $\varepsilon_N = 0.06$. This classifier was developed in Python3 using the OpenCV and NumPy libraries (https://github.com/heberlr/Image_Classifier).

Since our model is stochastic, we conducted a study to test the capability of our model to maintain the same classification. Thus, we executed 20 replicas of our model for the following parameters: $F_r = 50\%$, $b^* = 0.5$, and $T_p = 50h$. According to Figure 5, we found that 19 replicates were classified with plumes, 18 with cell escape, and all replicates without a necrotic nucleus. This experiment shows that the classification does not substantially change for the fixed parameters. After verifying this, we performed a study of the response associated with the parametric space (Figures 6-8).

#### Heterogeneous Oxygen Distribution

To investigate a scenario with multiple $O_2$ sources, we display the simulation of the model for the case in which we arranged oxygen sources randomly in the computational domain, adopting $F = 50$, $b = 0.5$, and $T_p = 50\ h$. This simulation showed that the distribution of oxygen modified the geometry of the tumor, changing the mechanical aspect of tumor growth. In Figure 9A, we found that the constant flow of GFP+ cells towards regions of greater oxygenation (proximity to blood vessel sections) implies a rapid saturation of cell density, losing invasive "plumes" and may even generate non-physical phenomena, such as the overlapping of cellular nuclei. Based on this observation, we added new mechanical restrictions on the dynamics of migration and proliferation. We applied restrictions based on the dimensionless mechanical pressures in each cell (`simple_pressure` of PhysiCell[1]). When the mechanical pressure is above a certain threshold as in regions of high cell density (10, estimated), the cell will not proliferate and will have a migration stimulus in the opposite direction. In Figure 9B, we present a simulation including negative feedback on proliferation and migration. Note that in this new formulation of the model, the invasive structures remain well defined along time evolution. Another interesting topic that could be included in this new approach would be incorporating the mechanical effects of cell intravasation into the blood vessels. Based on the mechanical pressure around the blood vessels, tumor cells would have a probability of breaking through the barrier to the basement membrane of the capillaries. This, in turn, would reduce the cell density near the vessels, allowing further proliferation and cell flux.

[1] https://github.com/MathCancer/PhysiCell/blob/master/documentation/User_Guide.pdf

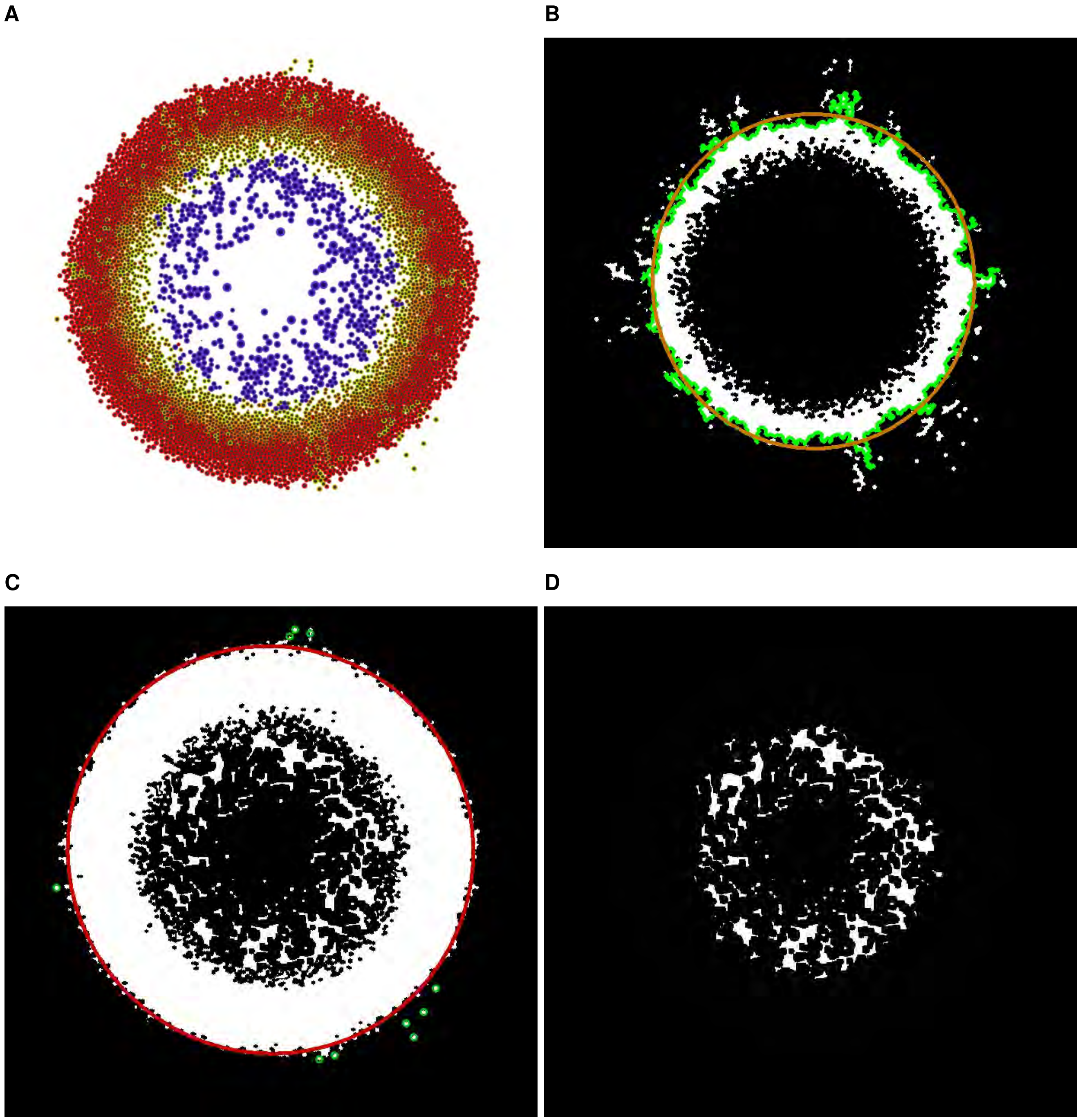


**Figure 4. Image classifier output.** (A) Output of the model using one simulation of the model at 100 hours with $F_r = 50\%$, $b^* = 0.5$, and $T_p = 50h$. (B) Image of the plumes Boolean test. The function $\mathcal{F}_{\mathcal{G}}$ in green and ellipse $\mathcal{E}_G$ in orange. (C) Image of the Boolean test for escaping cells. The ellipse $\mathcal{E}_T$ fitting the tumor in red, and the GFP+ cells escaping the tumor in green. (D) Image of necrotic area.

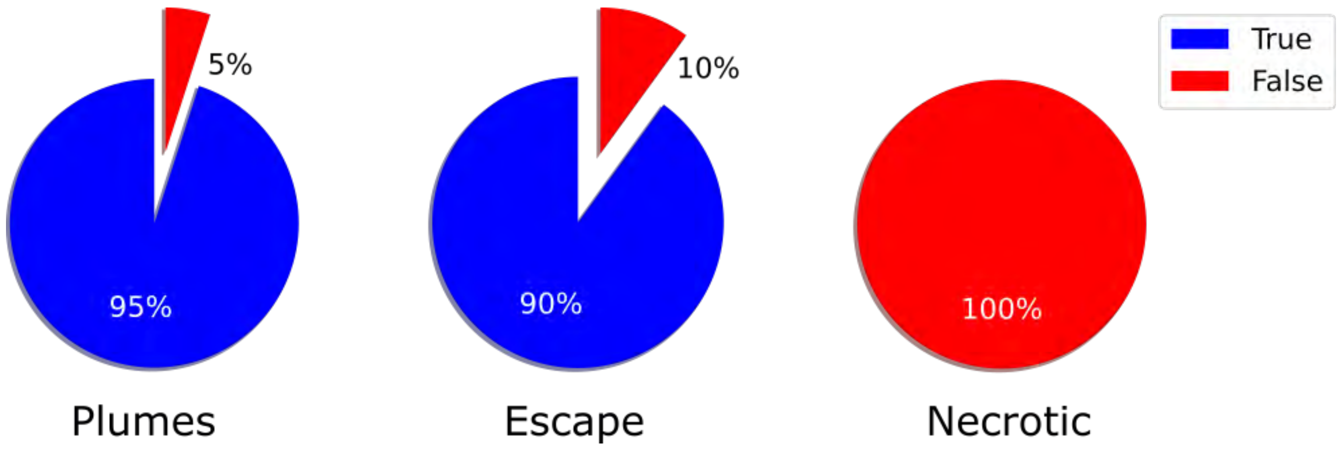


**Figure 5. Boolean image classification.** Percentage of the Boolean image classification of 20 replicates of the model (associated to 100h of tumor evolution), when $F_r = 50\%$, $b^* = 0.5$, and $T_p = 50h$.

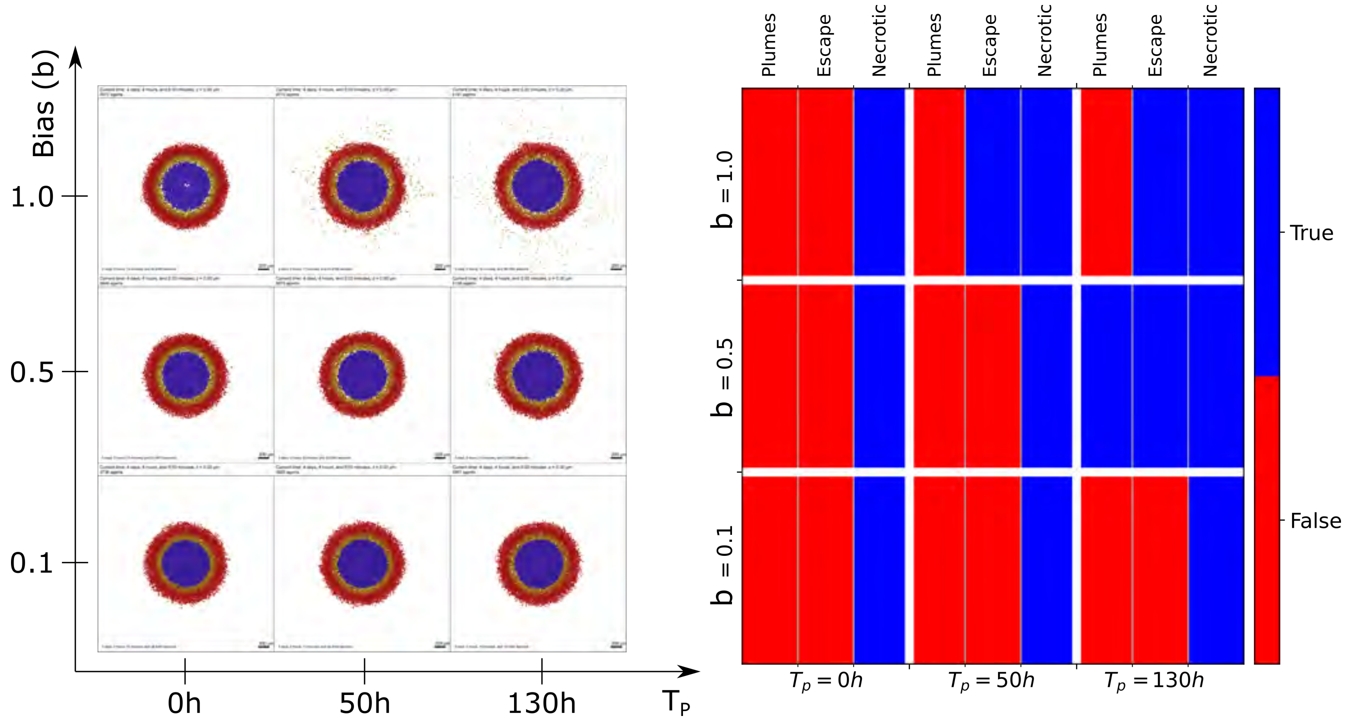


**Figure 6. Study varying b and $\mathbf{T_p}$ with $\mathbf{F_r = 10\%}$.** Images resultant from the proposed model (left) and associated Boolean classification (right). All simulations were evaluated at 100 hours after the initial condition.

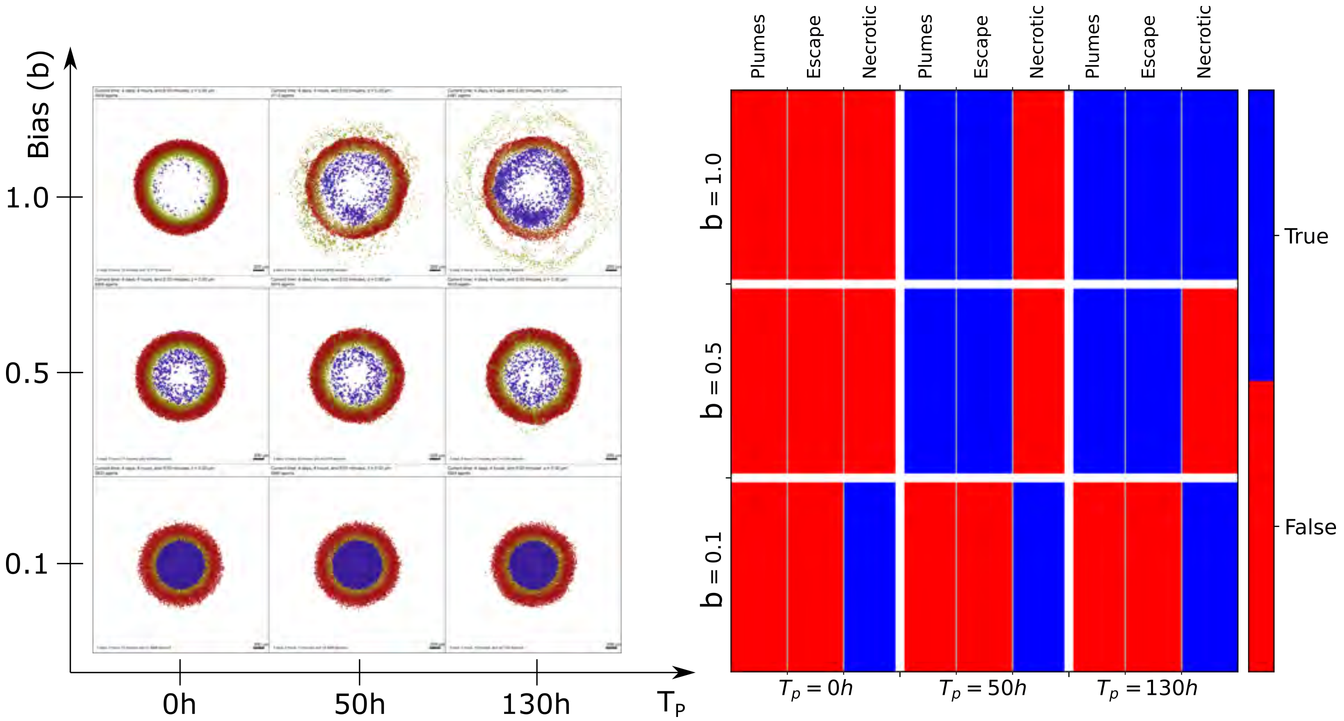


**Figure 7. Study varying b and $T_p$ with $F_r = 50\%$.** Images resultant from the proposed model (left) and associated Boolean classification (right). All simulations were evaluated at 100 hours after the initial condition.

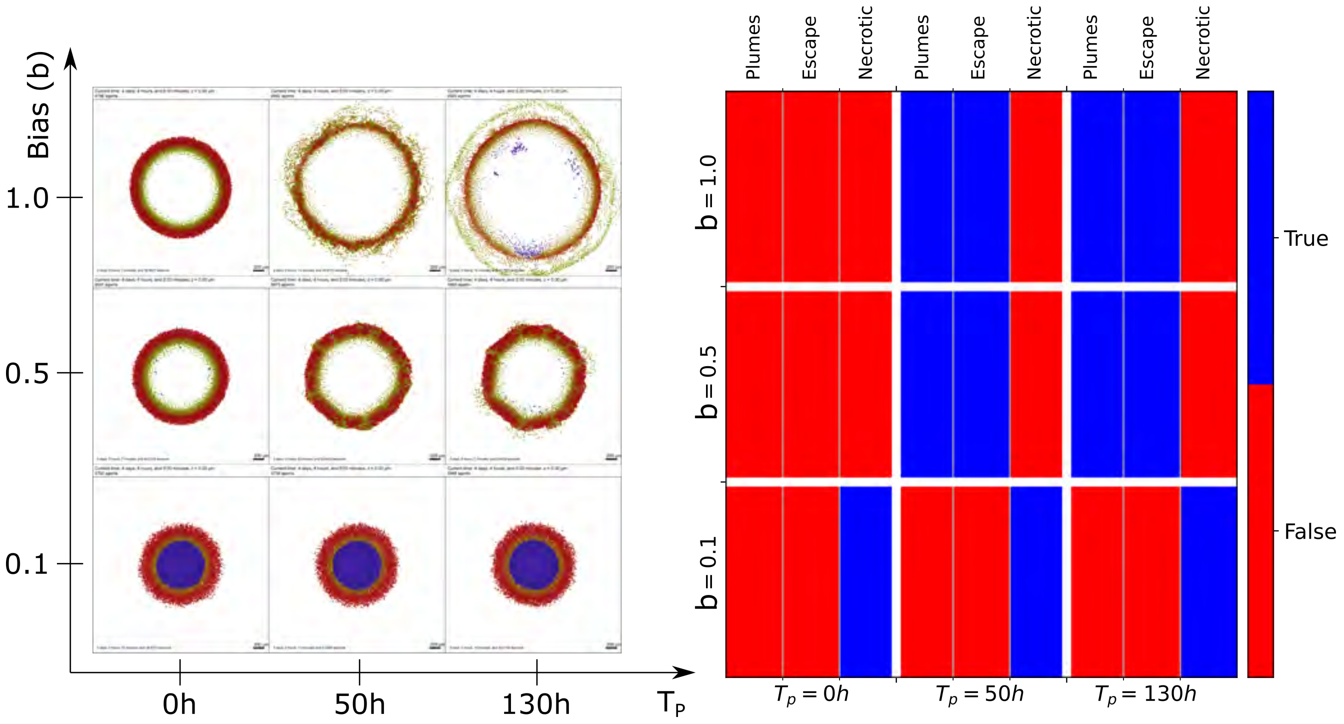


**Figure 8. Study varying b and $T_p$ with $F_r = 100\%$.** Images resultant from the proposed model (left) and associated Boolean classification (right). All simulations were evaluated at 100 hours after the initial condition.

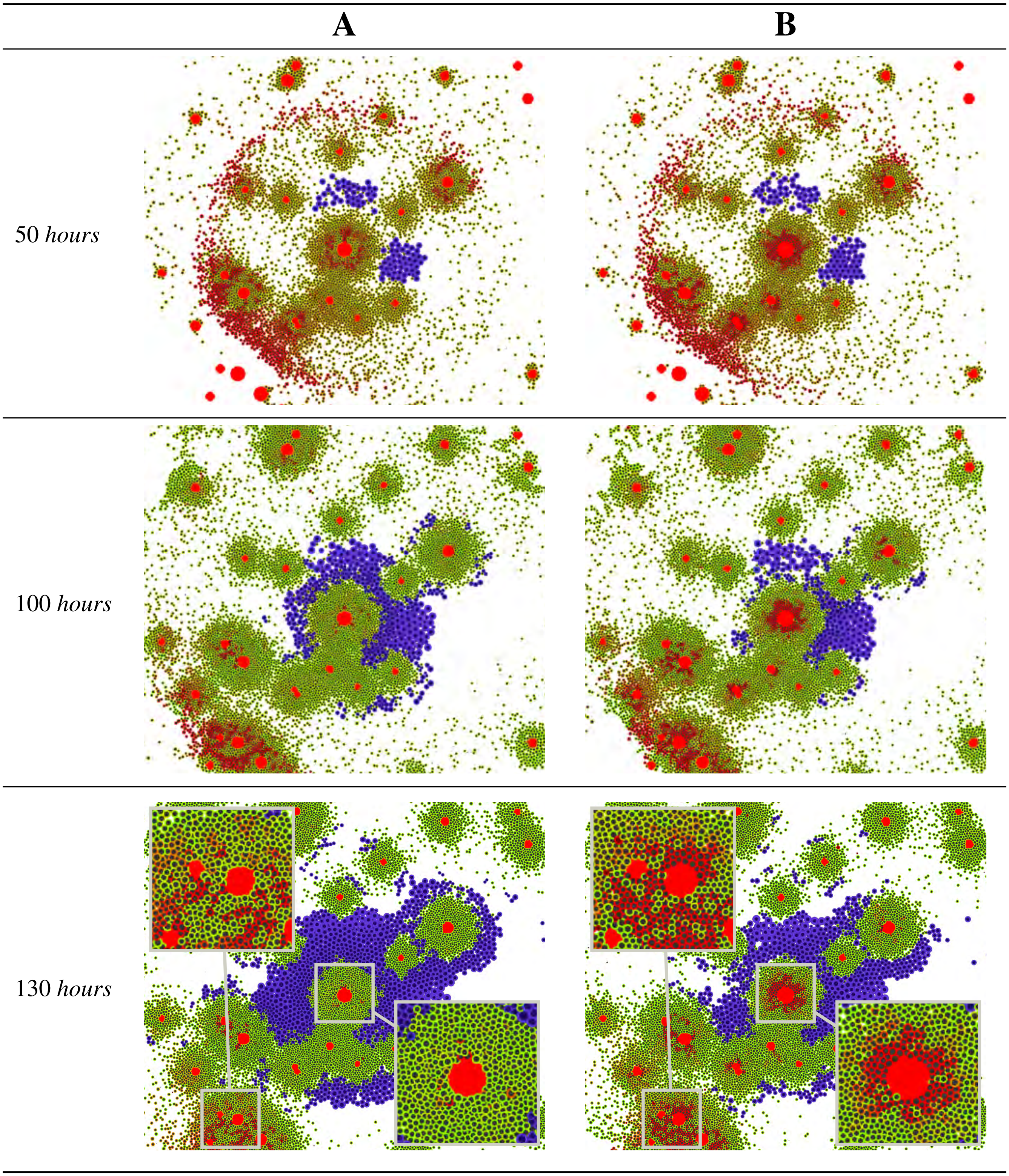


**Figure 9. Impact of heterogeneous oxygen distribution, $O_2$ sources randomly arranged (red circles).** (A) The computational model dissipate invasive structures and generate overlapping between cellular nuclei ($F = 50$, $b = 0.5$, and $T_p = 50$ $h$). (B) Mechanical feedback on proliferation and migration helps to maintain invasive structures.
